# NMD is required for timely cell fate transitions by fine-tuning gene expression and controlling translation

**DOI:** 10.1101/2020.07.07.180133

**Authors:** Elena Galimberti, Robert Sehlke, Michelle Huth, Marius Garmhausen, Merrit Romeike, Julia Ramesmayer, Sarah Stummer, Fabian Titz-Teixeira, Veronika Herzog, Anastasia Chugunova, Katrin Friederike Leesch, Laurenz Holcik, Klara Weipoltshammer, Laura Santini, Andreas Lackner, Arndt von Haeseler, Christa Bücker, Andrea Pauli, Christian Schoefer, Stefan L. Ameres, Austin Smith, Andreas Beyer, Martin Leeb

**Author notes:** equal contribution (Robert Sehlke, Michelle Huth).

## Abstract

Cell fate transitions depend on balanced rewiring of transcription and translation programmes to mediate ordered developmental progression. Components of the nonsense-mediated mRNA decay (NMD) pathway have been implicated in regulating embryonic stem cell (ESC) differentiation, but the exact mechanism is unclear. Here we show that NMD controls the translation initiation factor *Eif4a2* and its premature termination codon encoding isoform (*Eif4a2^PTC^*). NMD deficiency leads to translation of a specific truncated Eif4a2 protein, which elicits increased translation rates and causes significant delays in mouse ESC differentiation. Thereby a previously unknown feedback loop between NMD and translation initiation is established. Our results illustrate a clear hierarchy between KOs in severity of target deregulation and differentiation phenotype (*Smg5* > *Smg6* > *Smg7*), which highlights heterodimer-independent functions for Smg5 and Smg7. Together, our findings expose an intricate link between mRNA stability and translation initiation control that must be maintained for normal dynamics of cell state transitions.

## INTRODUCTION

Mouse embryonic stem cells (ESCs) capture the developmentally transient naïve pluripotent state *in vitro*. ESC self-renewal is maintained by an interactive transcription factor (TF)-network, which has been extensively characterized ^1, 2^. This network is decommissioned between embryonic day (E)4.5 and E5.5 in mouse development ^3^. Similar kinetics are maintained in ESCs, which irreversibly commit to differentiation 24-36 h after 2i withdrawal ^4–6^. Rex1 is a known marker of naïve pluripotency and high-resolution dissection of the exit from naïve pluripotency is facilitated by the availability of Rex1 promoter-driven destabilized GFP reporter ESC lines (Rex1::GFPd2) ^7, 8^. Rex1-GFP downregulation is initiated within 24 h after 2i withdrawal (N24) and completed after 48 h (N48). Extinction of Rex1-GFPd2 expression coincides with functional commitment to differentiation.

Several genome-wide screens have uncovered drivers of the exit from pluripotency ^9^, many of which are involved in transcriptional regulation and epigenetic modification. These screens also identified a large cohort of post-transcriptional regulators. RNA modifiers, such as m6A ^10, 11^; negative regulators of mRNA stability, such as Pum1 ^12^; and components of the nonsense-mediated mRNA decay (NMD) pathway ^12–14^ have been implicated in regulating ESC differentiation. Nonetheless, how post-transcriptional regulatory mechanisms contribute to cell fate changes remains poorly understood.

NMD is a translation coupled mechanism that promotes degradation of mRNAs containing a premature termination codon (PTC) ^15^. However, PTC-independent NMD activity has also been shown ^16–18^. NMD is triggered by phosphorylation of the RNA-helicase Upf1, which is essential for NMD. Degradation occurs either by Smg6-mediated endonucleolytic cleavage, or by exonucleolytic cleavage, mediated by a Smg5-Smg7 heterodimer ^15^. Transcriptome-wide analysis demonstrated that Smg6 and Smg5-Smg7 have highly overlapping mRNA targets ^19^. There is also evidence that Smg factors regulate telomere maintenance ^20–22^. Although NMD components constitute some of the top hits in genome-wide screens studying exit from naïve pluripotency, neither the contribution of individual NMD effector proteins, nor the key mRNA targets of NMD that lead to delayed cell fate transition are known.

Here we identify a role for NMD in ensuring normal differentiation kinetics by facilitating establishment of proper cell fate specific gene expression programmes and by controlling expression of Eif4a2, a key translation initiation factor. We identify the resulting Eif4a2-dependent increased translational activity in NMD-deficient ESCs as the main reason for their inability to properly commit to differentiation.

## RESULTS

### Variable degree of defects in exit from naïve pluripotency in NMD-deficient ESCs

To delineate the molecular function of NMD in the exit from naïve pluripotency, we generated Rex1-GFPd2 reporter ESC lines ^4, 23^ deficient for the three NMD downstream effectors *Smg5*, *Smg6* or *Smg7* (NMD KO ESCs), and corresponding rescue cell lines in which the missing NMD-factor was re-expressed (NMD rescue ESCs) (Extended Data Fig. 1a,b). NMD KO cells showed normal ESC morphology, cell cycle profile and telomere maintenance (Extended Data Fig. 1c,d), but exhibited pronounced delays in the exit from naïve pluripotency. This manifested in delayed downregulation of the Rex1-GFPd2 reporter 24 h after the onset of differentiation (N24) and delayed entry into commitment, assayed 72 h after 2i withdrawal (Fig. 1a,b). Both defects were restored in rescue cell lines, showing causality of NMD-factor deficiency for delays in exit from naïve pluripotency (Fig 1a,b). The three NMD-factor KOs displayed variable degrees of differentiation delays: the strongest effect was observed in the absence of *Smg5* and the weakest in the absence of *Smg7*. This does not align with the proposed strictly heterodimer-dependent function of Smg5 and Smg7, which would predict similar phenotypes for *Smg5* and *Smg7* KOs ^24–26^. The results therefore suggest a less strict heterodimer dependence than anticipated for the activity of either *Smg5* or *Smg7*.

**Fig. 1.**
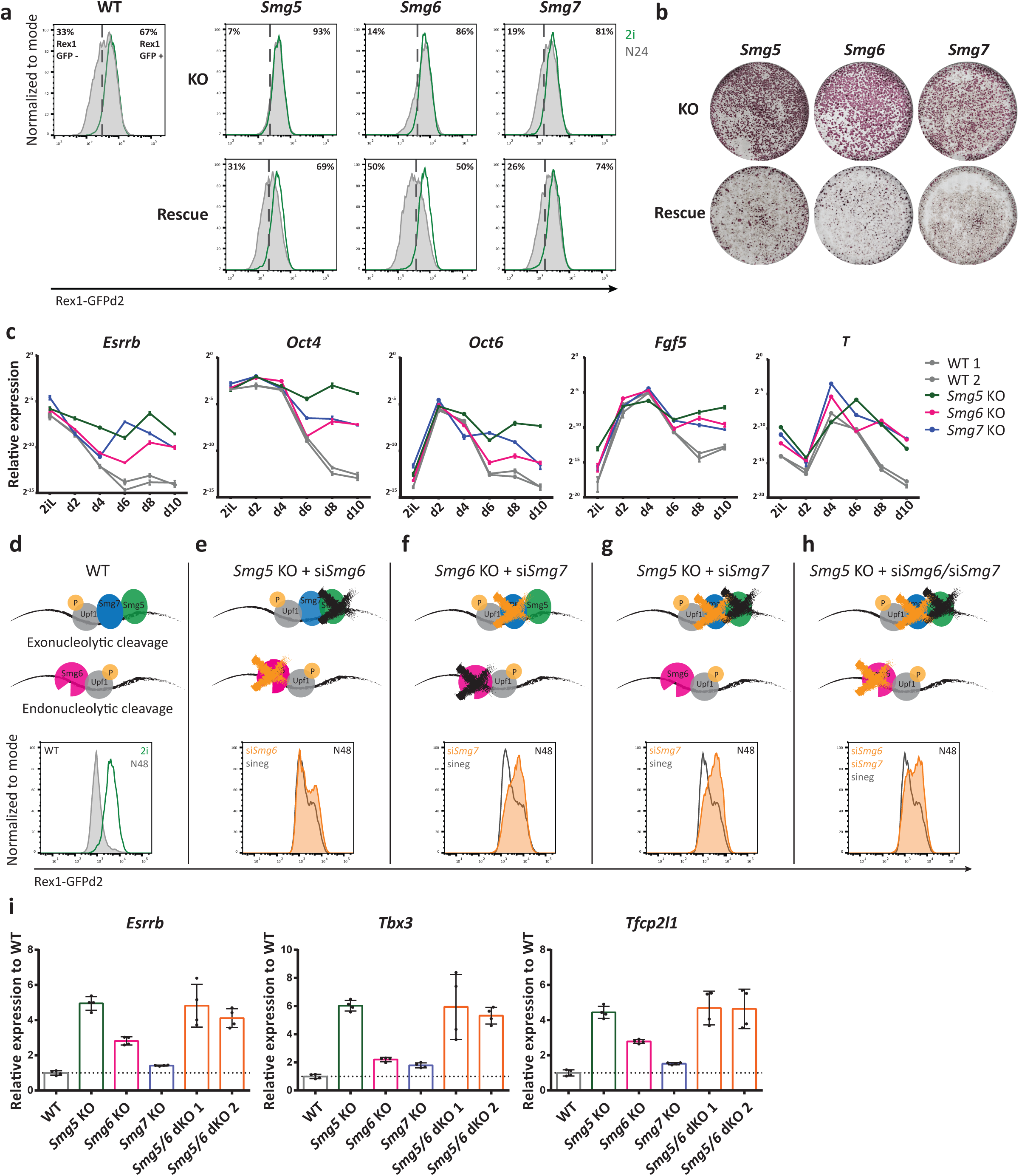
Hierarchy of defects in exit from naïve pluripotency in NMD-deficient ESCs. **a**, Rex1-GFPd2 analysis by FACS 24 h after induction of differentiation by withdrawing 2i in N2B27 medium (N24) in NMD KO, NMD rescue and WT ESCs. Green indicates profiles in 2i and grey indicates N24 profiles. **b**, Commitment assay in NMD KO and NMD rescue ESCs confirms commitment defects. Cells were re-plated in 2i medium after 72 h of differentiation. Only non-committed cells can reinitiate ESC self-renewal. Alkaline phosphatase staining was used to visualize ESC colonies. **c**, Representative experiment for expression kinetics of the indicated genes during a 10-day embryoid body (EB) differentiation assay. Mean and SD of technical replicates are plotted for each time point. Expression levels were normalized to 60S ribosomal protein L32 Rpl32 (*L32)*. **d-h**, Rex1-GFPd2 analysis at N48 of WT cells **(d)** and NMD KO and NMD dKO cells after siRNA KD of *Smg6* or *Smg7* (**e-h**). **i**, Expression of the indicated genes at N24 in NMD KO, *Smg5/Smg6* dKO and WT ESCs. Mean and SD are plotted for each cell line, n=2 biological replicates. Expression was normalized first to *Actin* and results are shown relative to WT. Unpaired t-test was used to calculate p-values (**** p < 0.02 unless indicated in the graphs).

To monitor the effect of NMD deficiency on long-term differentiation potential, we performed 3D aggregate embryoid body (EB) differentiation during a 10-day time-course. As expected, downregulation of naïve and primed pluripotency markers was severely impaired in NMD KO EBs. Although NMD KO aggregates upregulated the formative marker genes *Fgf5* and *Otx2* with similar kinetics to WT, the subsequent shutdown of the formative program, observed in WT cells between day 4 (d4) and d6, was impaired (Fig. 1c). Furthermore, expression of the endo-mesoderm defining TF *Brachyury* (*T*) was not properly downregulated after a peak of expression at d4 to d6, suggesting a general function of NMD in shaping transcriptomes during early cell fate decisions (Fig. 1c). Further supporting a role for NMD in regulating differentiation, teratomas derived from *Smg6* KO ESCs showed a lower degree of differentiation than WT and rescue controls (Extended Data Fig. 1e). Together, these data show a global differentiation defect of Smg factor-deficient cells, with a variable degree of phenotypic strengths (Smg5 > Smg6 > Smg7).

### Cooperativity between NMD factors regulates NMD and exit from naïve pluripotency

To investigate potential cooperativity between NMD effectors, we performed siRNA-mediated knockdowns of *Smg6* and *Smg7* in *Smg5* and *Smg6* KO ESCs (Fig. 1d-h and Extended Data Fig. 1f). Thereby, we generated cells double- and triple-depleted for NMD downstream effectors. WT cells showed a near complete loss of Rex1-GFPd2 expression at N48 (Fig. 1d). siRNA-mediated depletion of *Smg6* in *Smg5* KO cells resulted in only a minor, non-synergistic increase in the differentiation delay assessed in Rex1-GFPd2 reporter assays (Fig. 1e), despite the strong defects observed in both single KOs. By contrast, the depletion of *Smg7* in a *Smg6* KO background yielded a strong synergistic differentiation phenotype (Fig. 1f). Similarly, co-depletion of both heterodimer members, *Smg5* and *Smg7*, exhibited a clear synergistic effect (Fig. 1g), further highlighting heterodimer-independent functions for Smg5 and Smg7. Combined depletion of all the NMD effectors by double knockdown of *Smg6* and *Smg7* in *Smg5* KO ESCs resulted in differentiation delays on par with *Smg6/7* or *Smg5/7* double depletion (Fig. 1h). To assess the impact of NMD effector depletion and co-depletion on NMD-specific mRNA target gene expression, we assessed levels of the known NMD target gene *Gadd45b* by RT-qPCR ^16^. Consistent with observed differentiation delays, we observed strong genetic interactions between *Smg6* and *Smg7* as well as between *Smg5* and *Smg7*, and weaker interactions between *Smg5* and *Smg6* regarding *Gadd45b* transcript abundance (Extended Data Fig. 1g). Together, this suggests that differentiation delays scale with the extent of NMD impairment.

To further study the cooperativity of NMD factors we sought to generate stable NMD double KO (dKO) ESCs. However, attempts using CRISPR/Cas9 to generate all possible dKO ESC lines yielded only *Smg5/Smg6* (*Smg5/6*) dKO ESCs. Neither *Smg5/Smg7* nor *Smg6/Smg7* dKOs could be established, despite multiple attempts using efficient gRNAs. This suggests an essential role of *Smg7* in ESCs in the absence of its heterodimerization partner *Smg5* or in the absence of *Smg6*. *Smg5/6* dKO cells showed a deficiency in downregulating naïve TF mRNAs similar to that observed in *Smg5* single KO cells, indicating a dominant role of *Smg5* in regulating differentiation-relevant RNA-homeostasis programmes (Fig. 1i).

Taken together, *Smg5*, *Smg6* and *Smg7* KO ESCs exhibit variable degrees of differentiation defects and NMD downstream factors act synergistically during the exit from pluripotency. Smg7 possesses a central, facultatively essential function: It can be depleted without resulting in strong effects on early differentiation, but an essential role is revealed in the absence of either *Smg5* or *Smg6*.

### Smg7 is necessary and sufficient for pUpf1 binding, independently of Smg5

Continuous NMD activity relies on cyclic phosphorylation and dephosphorylation of Upf1. We observed increased pUpf1 levels in all NMD KO cells (Fig. 2a), in line with previous reports ^25^. Notably, pUpf1 levels followed the same trend as the differentiation phenotype and deregulation of the NMD target *Gadd45b* (pUpf1 levels in Smg5 KO > Smg6 KO > Smg7 KO > WT ESCs). Upf1 is phosphorylated by the kinase Smg1 ^27, 28^. *Smg1* mRNA is itself an NMD target, but in contrast to pUpf1 is upregulated to a similar extent in all three NMD KO ESCs (Extended Data Fig. 2a). Therefore, graded increase of pUpf1 levels in NMD KOs is unlikely to be a direct effect of higher *Smg1* expression. Impaired Upf1 dephosphorylation may therefore underlie increased pUpf1 levels in NMD mutant cells. The three NMD effectors Smg5, Smg6 and Smg7 have all been implicated in the recruitment of PP2A to dephosphorylate Upf1 ^25, 26^, but the strongest increase of pUpf1 in Smg5 KO ESCs suggests that Smg5 acts as the main PP2A recruiter during NMD (Extended Data Fig. 2b).

**Fig. 2.**
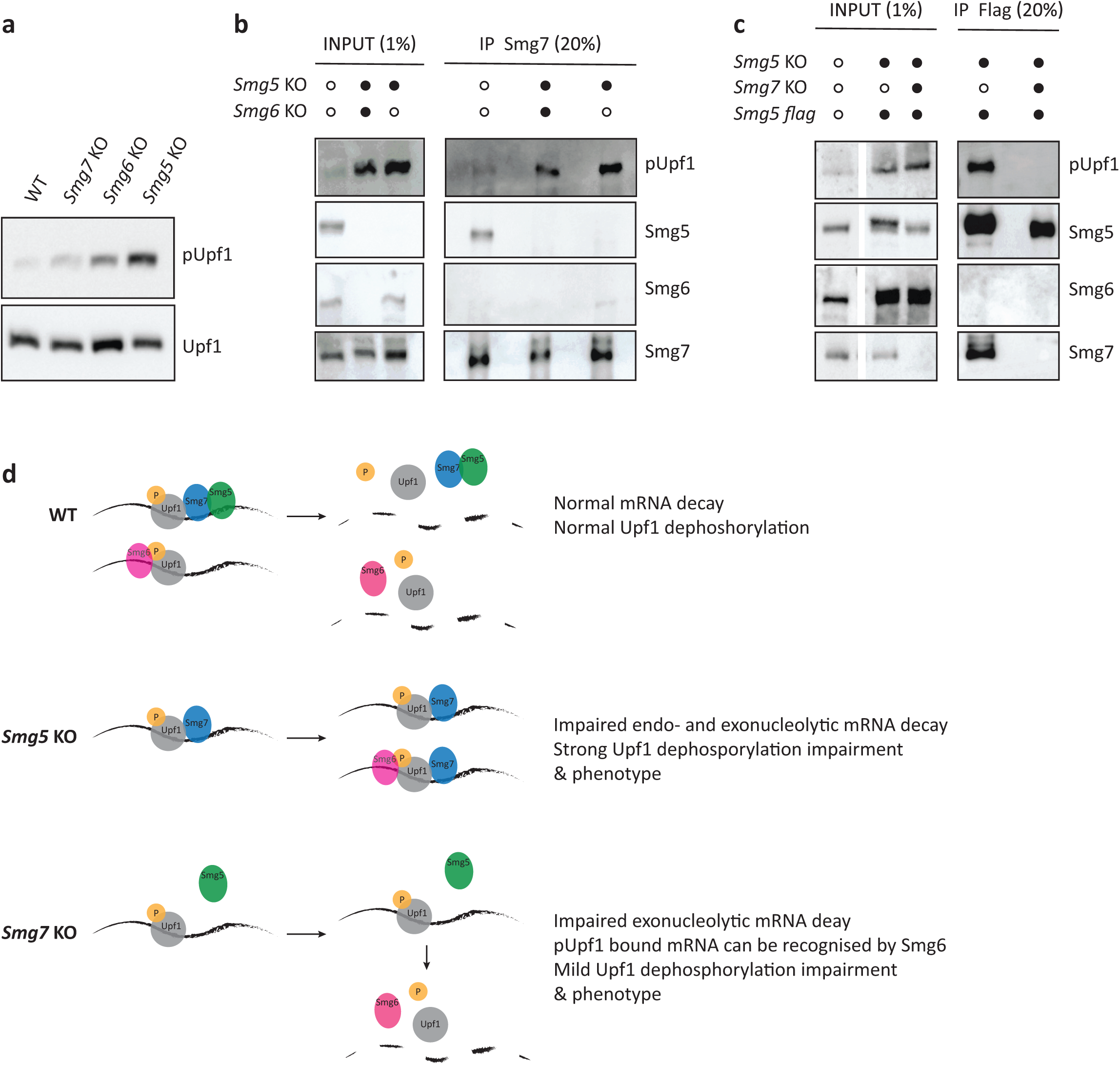
Smg7 is necessary and sufficient for pUpf1 binding, independently of Smg5. **a**, Western blot analysis for Upf1 and pUpf1 expression in NMD KO and WT ESCs. **b**, Western blot showing results of Smg7 co-IP in the indicated cell lines after cyanase treatment for RNA removal. Antibodies used are indicated in the figure. **c**, Western blot analysis of Smg5 co-IP (flag antibody co-IP in Smg5KO^Smg5flagrescue^ cells) in the indicated cell lines after cyanase treatment. Antibodies used are indicated in the figure. **d**, Schematic illustration of the proposed interactions between NMD effectors and pUpf1 dephosphorylation in WT, *Smg5* and *Smg7* KO ESCs that lead to a graded deficiency in NMD and exit from naïve pluripotency.

To identify the molecular basis for an apparent heterodimer-independent function of *Smg5* and *Smg7* and for the observed genetic interactions between Smg factors, we performed co-immunoprecipitation (co-IP) experiments. We precipitated Smg5 or Smg7 in WT and NMD KO ESCs, in the presence and absence of RNA. Smg5 and pUpf1, but not Smg6, co-immunoprecipitated with Smg7 in an RNA-independent manner in WT ESCs (Fig. 2b). Interestingly, Smg7 efficiently bound pUpf1 in the absence of its heterodimerization partner Smg5 (Fig. 2b). The Smg7-pUpf1 interaction was also maintained in *Smg5/6* dKO ESCs, suggesting that Smg7 can interact with pUpf1 independently of other downstream NMD effectors. Conversely, however, Smg5 failed to bind pUfp1 in the absence of *Smg7* (Fig. 2c), indicating that Smg7 is the critical docking site for pUpf1 during NMD and that pUpf1 binding to Smg7 is independent of Smg5. When we performed the same co-IP experiments in the presence of RNA, we observed an interaction between Smg5 or Smg7 and Smg6 in the absence of the respective heterodimerization partner (Extended Data Fig. 2c,d). This RNA-dependent interaction was not observed in WT (Fig. 2b,c).

Taken together, biochemical analysis and viability of *Smg5/6* dKO ESCs indicate that Smg7 can bind to pUpf1 in the absence of Smg5 to sustain limited NMD function. Smg5, however, is unable to interact with pUpf1 in the absence of Smg7 (Fig. 2c and Extended Data Fig. 2d). The strong differentiation phenotype of *Smg5*-deficient cells can be explained by efficient binding of pUpf1 by Smg7 alone and subsequent inefficient dephosphorylation and target degradation, resulting in a ‘poisoning’ of both branches of NMD (Fig. 2d). By contrast, deficiency in Smg7 results in failure to recruit components of the exonucleolytic branch, but allows potential recognition and degradation by Smg6, resulting in only a mild NMD target deregulation and differentiation phenotype.

### Integrating transcriptome-wide gene expression with mRNA half-life analyses identifies relevant NMD targets during the exit from naïve pluripotency

To uncover the molecular mode of action by which NMD regulates the exit from naïve pluripotency, we applied a combinatorial approach based on the following logic: an NMD target transcript relevant in the context of differentiation must show elevated expression levels, following the same graded pattern as the differentiation phenotype (Smg5 > Smg6 > Smg7) and a concomitant increased mRNA half-life.

To this end, we first performed RNA-seq of *Smg5* KO, *Smg6* KO, *Smg7* KO and WT cells in 2i and after 24 h of differentiation (N24) to assess the impact of NMD factor depletion on global gene expression (Fig. 3a - d and Extended Data Table 1). In 2i, we identified 881 transcripts deregulated in NMD KO ESCs compared to WT (adj. p ≤ 0.01), 516 of which were significantly upregulated and 252 significantly downregulated in all three NMD KO ESCs. This shows a strong overlap in deregulated genes between all three KO cell lines; only 113 transcripts did not show the same directionality in transcriptional changes between all NMD KO cells (Fig. 3a). Overall, upregulated transcripts (clusters 2 and 4), showed a graded increase in deregulation (*Smg5* KO > *Smg6* KO > *Smg7* KO), similar to the KO phenotypes (Fig. 3b). 256 out of 564 transcripts belonging to clusters 2 and 4 were significantly upregulated in all three NMD KO cell lines and showed a graded response to Smg-factor depletion, suggesting that genes found within this group are responsible for the observed differentiation delays.

**Fig. 3.**
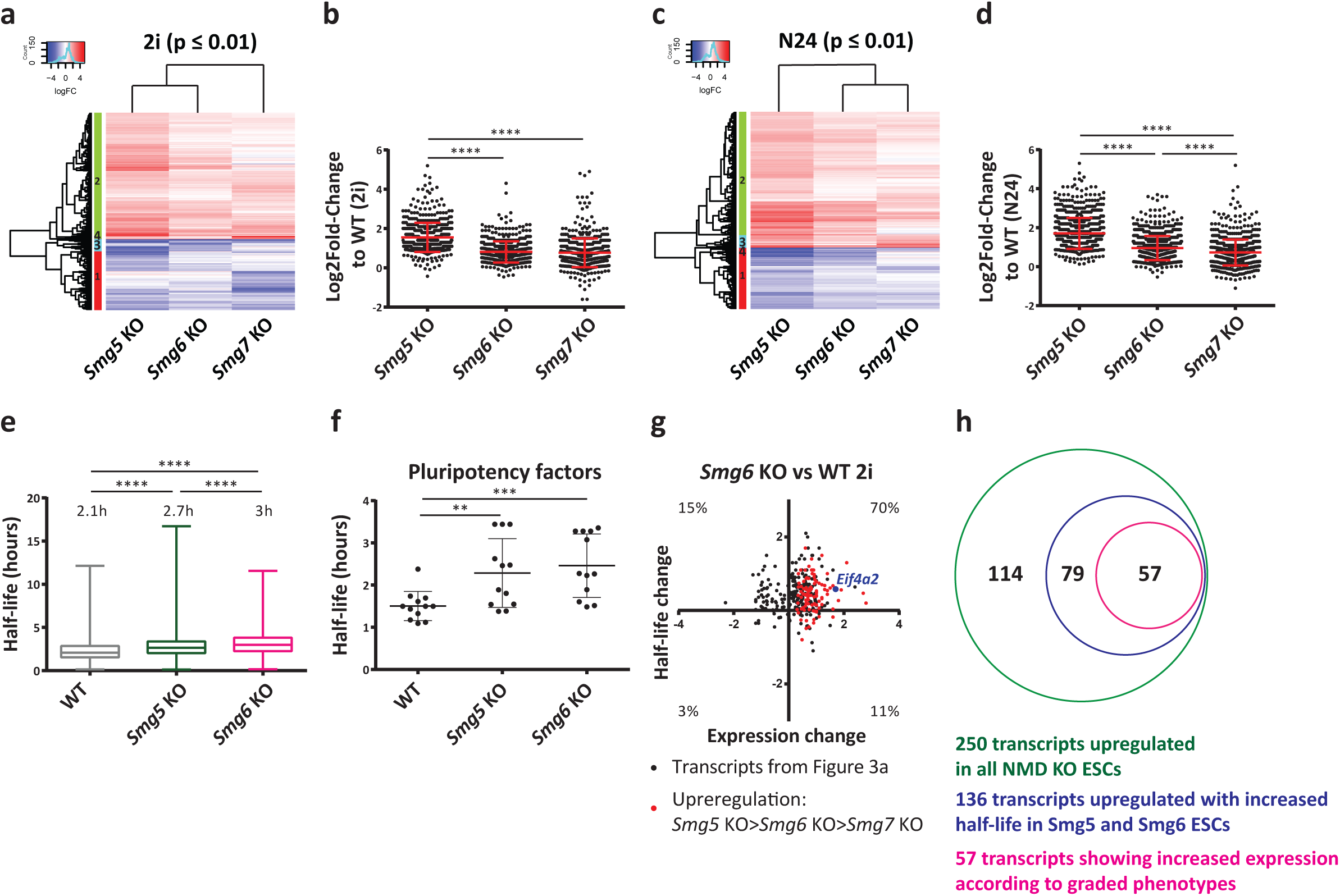
Integrating transcriptome-wide gene expression with mRNA half-life analyses identifies relevant NMD targets during the exit from naïve pluripotency. **a**, Differentially expressed genes between NMD KO and WT ESCs in self-renewal (2i) (p-adj. ≤ 0.01). p-adj. values were calculated testing against the null hypothesis of a fold change < |1.5|. **b**, Log2fold change of deregulated transcripts in 2i belonging to clusters 2 and 4 (564 transcripts). Mean and SD are indicated in the graph. Unpaired t-test was used to calculate p-values. (**** p-value < 0.0001). **c**, Differentially expressed genes between NMD KO and WT ESCs at N24 (p-adj. ≤0.01). p-adj. values were calculated testing against the null hypothesis of a fold change < |1.5|. **d**, Log2fold change of deregulated transcripts at N24 belonging to clusters 2 and 3 (808 transcripts). Mean and SD are indicated in the graph. Unpaired t-test was used to calculate p-values. (**** p-value < 0.0001). **e**, Half-life analysis in WT, *Smg5* KO and *Smg6* KO ESCs. Half-life was calculated in hours based on a pulse chase time course. Mean and SD are indicated in the graph. Paired t-test was used to calculate p-values (**** p-value < 0.0001). **f**, Half-life of pluripotency factors (*Esrrb, Klf2, Klf5, Nanog, Pou5f1, Sall4, Sox2, Stat3, Tbx3*) in WT, *Smg5* KO and *Smg6* KO ESCs. Half-lives were calculated in hours. Mean and SD are indicated in the graph. Paired t-test was used to calculate p-values (** p-value = 0.006, *** p-value = 0.0006). **g**, Transcriptome-wide comparison of changes in mRNA expression and half-life in *Smg6* KO. Half-life changes between WT and Smg6 KO for the 321 out of 516 transcripts upregulated in NMD KO ESCs for which half-lives could be calculated were calculated (see Figure 3a). 150 of these 321 transcripts showed an increase in half-life (log2FC ≥ 0.2) in both Smg5 and Smg6 KOs. Transcripts behaving in accordance with the graded phenotype (Smg5 > Smg6 > Smg7) are depicted in red. **h**, Schematic illustration of the approach used to identify NMD targets relevant for the differentiation phenotype in ESCs. First significantly upregulated transcripts (see Figure 3a) with measurable half-lives in WT and Smg KOs were selected (250). 136 transcripts showed a concomitant increase in half-life in Smg5 and Smg6 KOs (log2FC ≥ 0.2). 57 of these transcripts were upregulated according to the graded phenotypes.

GO analysis of the 256 graded upregulated transcripts in 2i revealed an enrichment for methyltransferase and helicase activity. The remaining upregulated transcripts in 2i showed enrichment for telomere maintenance and helicase activity. Within downregulated genes we detected an enrichment for transcripts involved in neural development, somite development and other differentiation related processes (Extended Data Fig. 3a and Extended Data Table 2). This suggests that leaky expression of the differentiation programme in 2i is dampened in NMD-deficient cells.

At N24, we identified 1,174 deregulated transcripts in NMD KO cells compared to WT cells (adj. p ≤ 0.01). Among these, 727 were upregulated in all NMD KO cells and 336 were downregulated in all NMD KO cells (Fig. 3c). Out of the 808 upregulated transcripts belonging to clusters 2 and 3, 434 transcripts were upregulated following the graded pattern (*Smg5* KO > *Smg6* KO > *Smg7* KO) (Fig. 3d). GO analysis of these 434 transcripts revealed enrichment for pluripotency-related terms, such as LIF response and stem cell maintenance (Extended Data Fig. 3b and Extended Data Table 2). This was not evident for the 293 transcripts not showing graded upregulation. Within downregulated transcripts, we detected an enrichment of differentiation-related terms, such as neural tube development and pattern specification (Extended Data Fig. 3b and Extended Data Table 2). This suggested that genes showing graded expression are involved in and reflect the different levels of differentiation delays seen in NMD KO cells.

Previous reports showed increased c-Myc levels in NMD-deficient ESCs and proposed a causative involvement of c-Myc in NMD KO induced differentiation delays ^13^. However, we did not detect increased c-Myc transcript levels by RNA-seq or protein levels on western blots in any of the three NMD KO ESCs (Extended Data Fig. 3c,d). On the contrary, we reproducibly detected reduced levels of c-Myc upon NMD disruption. Furthermore, genetic depletion of *c-Myc* in NMD KO ESCs did not rescue the differentiation delays (Extended Data Fig. 3e,f) indicating that c-Myc is not relevant for the differentiation defects observed in NMD KO ESCs.

Elevated transcript levels of direct NMD targets are expected to result from increased half-lives, based on impairment of mRNA degradation. Therefore, we used SLAM-seq to measure transcriptome-wide half-lives ^29^. We detected a global increase of transcript half-lives in *Smg5* and *Smg6* KO ESCs compared to WT ESCs (Fig. 3e). We were able to calculate half-lives for 4,342 transcripts; 3,062 of which exhibited a longer half-life in NMD KO cells (Fig. 3e). On average, mRNA half-life was significantly longer in NMD KOs than in WT (p < 10^-4^), increasing from 2.1 h in WT to 2.7 and 3 h in *Smg5* KO and *Smg6* KO, respectively (Fig. 3e). As a cohort, pluripotency TF-encoding mRNAs exhibited a significant increase in half-lives (Fig. 3f) similar to the overall increase of half-lives across the entire transcriptome. Both identity of transcripts and amplitude of half-life changes showed a strong overlap between *Smg5* and *Smg6* KO ESCs (Extended Data Fig. 3g). Transcriptome-wide comparisons showed that the vast majority of transcriptionally upregulated genes showed a concomitant increase in half-lives (Fig. 3g and Extended Data Fig. 3h).

To identify bona-fide NMD targets with relevance for the differentiation delay phenotype, we investigated a group of 250 genes that were significantly upregulated in all three *Smg* KO ESCs in 2i for which we could also calculate half-lives. Of these, 136 showed an increase in half-lives in both *Smg5* and *Smg6* KOs, and 57 showed an expression pattern in accord with the degree of phenotype-strength (Fig. 3h, Extended Data Fig. 3i and Extended Data Table 3). These 57 genes were involved in various cellular processes, such as tRNA modification, TOR-signaling and calcium transport (Extended Data Table 3). None of the naïve pluripotency factors was part of this group, suggesting that direct regulation of naïve pluripotency factors by NMD is not the primary cause for the observed differentiation delays in NMD KO ESCs. Taken together, we identified a cohort of 57 direct NMD target genes, with an expression pattern suggestive of an involvement in controlling the exit from naïve pluripotency.

### Eif4a2 is a bona-fide NMD target in ESCs and its NMD sensitivity is evolutionarily conserved

Among the 57 NMD targets with potential phenotypic relevance, our attention was drawn to the RNA helicase *Eif4a2,* a component of the Eif4F translation initiation complex, which mediates 5’ cap recognition by the small ribosomal subunit ^30, 31^. Several layers of evidence suggest *Eif4a2* as a bona-fide NMD target. Firstly, the *Eif4a2* locus encodes for two splice-isoforms: one full-length protein, and one PTC-containing isoform (Fig. 4a). Secondly, *Eif4a2* mRNA levels were significantly increased after translation inhibition by cycloheximide (CHX) treatment, a hallmark of bona-fide NMD targets (Fig. 4b). CHX response was similar to that observed for *Gadd45b*. The PTC-isoform (*Eif4a2^PTC^*) showed an even more pronounced sensitivity to CHX treatment. Thirdly, *Eif4a2^PTC^* transcript levels were upregulated in a graded fashion (*Smg5* KO > *Smg6* KO > *Smg7* KO) throughout a long-term EB differentiation time-course, consistent with phenotypic relevance (Fig. 4c). Full length Eif4a2 (Eif4a2^FL^) protein levels were increased in NMD KO cells (Fig. 4d). The *Eif4a2^PTC^* isoform produced a protein that was weakly detected by western blot analysis in WT and NMD rescue cells and strongly increased in all three NMD KO ESCs, both in 2i and at N24 (Fig. 4d and Extended Data Fig. 4a). This indicates that while expression levels are tightly regulated by NMD, the Eif4a2^PTC^ protein shows low levels of expression in WT cells. The upregulation of *Eif4a2* was unique among Eif4F complex members, since the others, including the close homologue Eif4a1, showed neither increased expression nor significantly longer half-lives in NMD KOs (Fig. 4d, Extended Data Fig. 4b,c and Table S3), nor did they react to CHX treatment (Extended Data Fig. 4d). Taken together, this identifies *Eif4a2* as a bona-fide NMD target in ESCs and shows that NMD disruption leads to the production of an *Eif4a2^PTC^* protein.

**Fig. 4.**
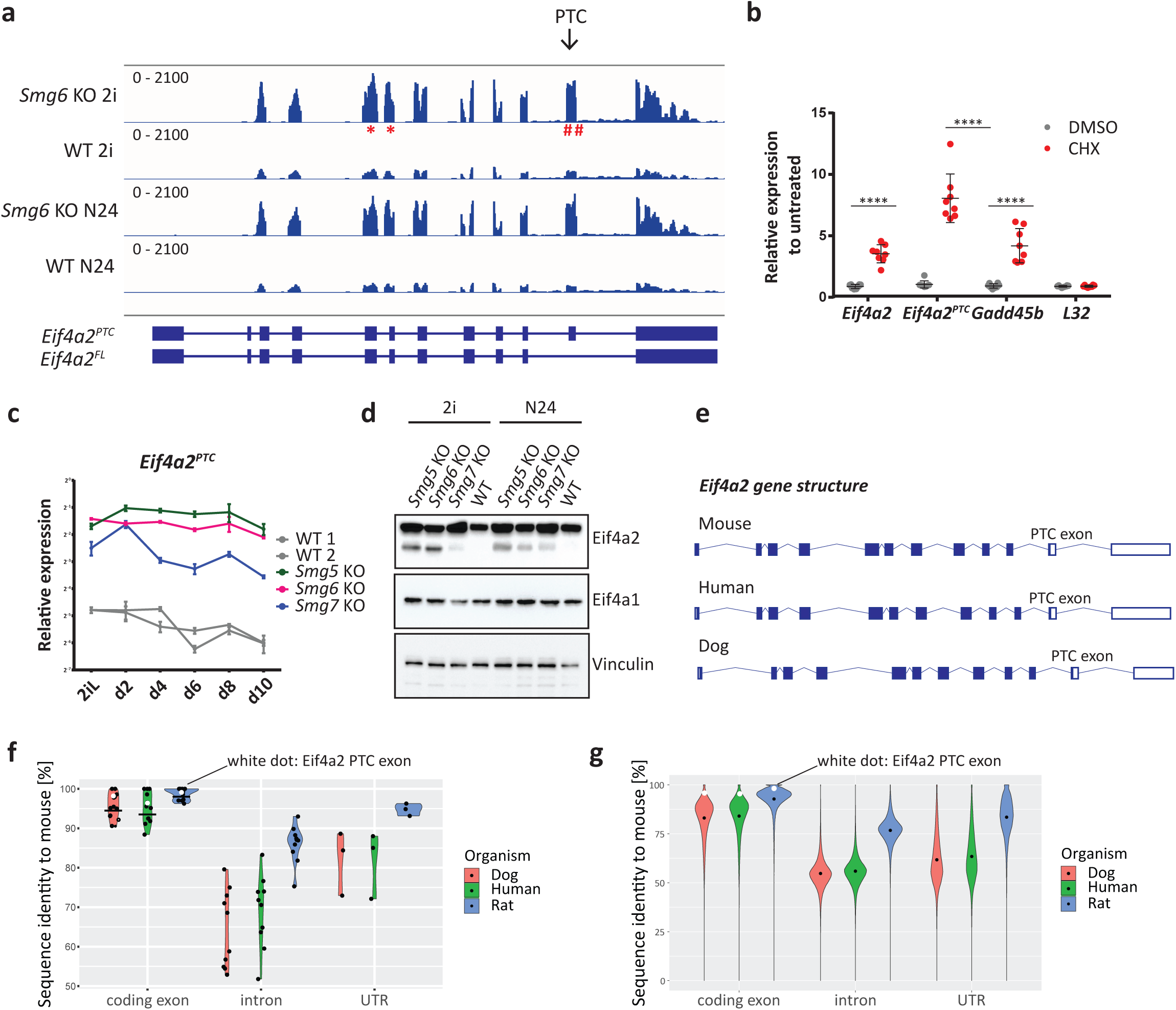
Eif4a2 is a bona-fide NMD target in ESCs. **a**, Genome browser view of the *Eif4a2* locus visualizing RNA-seq results in *Smg6* KO and WT ESCs in 2i and at N24. * indicates qPCR primers amplifying all *Eif4a2* isoforms. # indicates qPCR primers amplifying the *Eif4a2^PTC^* isoform. **b**, Expression levels of the indicated genes after treatment with CHX or DMSO in WT ESCs. Mean and SD are plotted, n=4 biological replicates. Unpaired t-test was used to calculate p-values (**** p < 0.0001). **c**, Representative experiment for expression kinetics of the indicated genes during an EB differentiation assay as in Figure 1c. Mean and SD from technical replicates are plotted for each time point. Expression levels were normalized to *L32*. **d**, Western blot analysis of Eif4a2 and Eif4a1 protein levels in NMD KO and WT ESCs in self-renewal (2i) and at N24. Vinculin was used as loading control. **e**, Schematic representation of the gene structure of Eif4a2 in mouse, human and dog. **f**, Percentage of sequence identity to mouse of Eif4a2 between dog, rat and human. White dots indicate the PTC-containing exon of *Eif4a2* and black lines indicate the mean. **g**, Percentage of sequence identity to mouse of the transcriptome between dog, rat and human. White asterisks indicate the PTC-containing exon of *Eif4a2* and the black dots indicate the mean.

The linkage of NMD to *Eif4a2* expression appears to be conserved in evolution. The gene structure of *Eif4a2* is strikingly similar between mouse, dog and human (Fig. 4e). The mouse *Eif4a2* coding sequence showed identities of 94%, 95% and 98.5% to human, dog and rat, respectively (Fig. 4f). Despite the PTC-exon being under no apparent selective pressure to maintain protein-coding potential (PTC at nucleotide position 8 out of 107 in this exon), we observed 96.4%, 98.2% and 99% nucleotide identity for human, dog and rat, respectively, on par with the other well conserved *Eif4a2* exons. Conservation of UTRs and introns was significantly lower (Fig. 4f). In a transcriptome-wide comparison, the *Eif4a2* PTC exon showed high conservation at the nucleotide level: less than 5% of all exons and less than 3% of all UTRs showed higher conservation rates between mouse and rat than the *Eif4a2^PTC^* exon (Fig. 4g and Extended Data Fig. 4e-g). In the comparisons with human and dog these percentages were even lower. Therefore, the potential of Eif4a2 to be regulated by NMD is a feature conserved in mammalian evolution.

### *Eif4a2* is causative for defects in exit from naïve pluripotency in NMD-deficient ESCs

To delineate a potential causative relationship between increased *Eif4a2* levels and the differentiation defect observed in NMD-deficient cells, we generated *Eif4a2* KO cells and NMD (*Smg5* or *Smg6*) / *Eif4a2* double deficient cells (collectively referred to as NMD/*Eif4a2* dKO) by deleting all potential *Eif4a2* isoforms or specifically *Eif4a2^PTC^* (Fig. 5a and Extended Data Fig. 5a). All *Eif4a2* KO cell lines proliferated normally, showed normal ESC morphology and could be maintained for more than 10 passages. Deletion of the PTC exon resulted in an increase in full length Eif4a2 protein levels in WT and NMD KO ESCs (Fig. 5a). Both the complete absence of *Eif4a2* or specific deletion of *Eif4a2^PTC^* accelerated Rex1-GFPd2 downregulation kinetics in WT cells (Fig. 5b), suggesting that increased expression of *Eif4a2^PTC^* and not full-length Eif4a2 is the major cause of the differentiation block. In NMD/*Eif4a2^FL^* and NMD/*Eif4a2^PTC^* dKO cells, differentiation kinetics were substantially rescued in a Rex1-GFPd2 assay (Fig. 5b). Accordingly, *Klf4* and *Nanog* expression levels were downregulated to near WT-levels at N30 (Fig. 5c). Similar effects were observed for protein levels of the naïve TFs Nanog and Esrrb at N24 (Extended Data Fig. 5b). Phenotypic rescue in NMD/*Eif4a2^PTC^* double KO cells, in which Eif4a2^FL^ protein levels are even further increased, supports the proposition that Eif4a2^PTC^ is the major mediator of the observed defect in exit from naïve pluripotency.

**Fig. 5.**
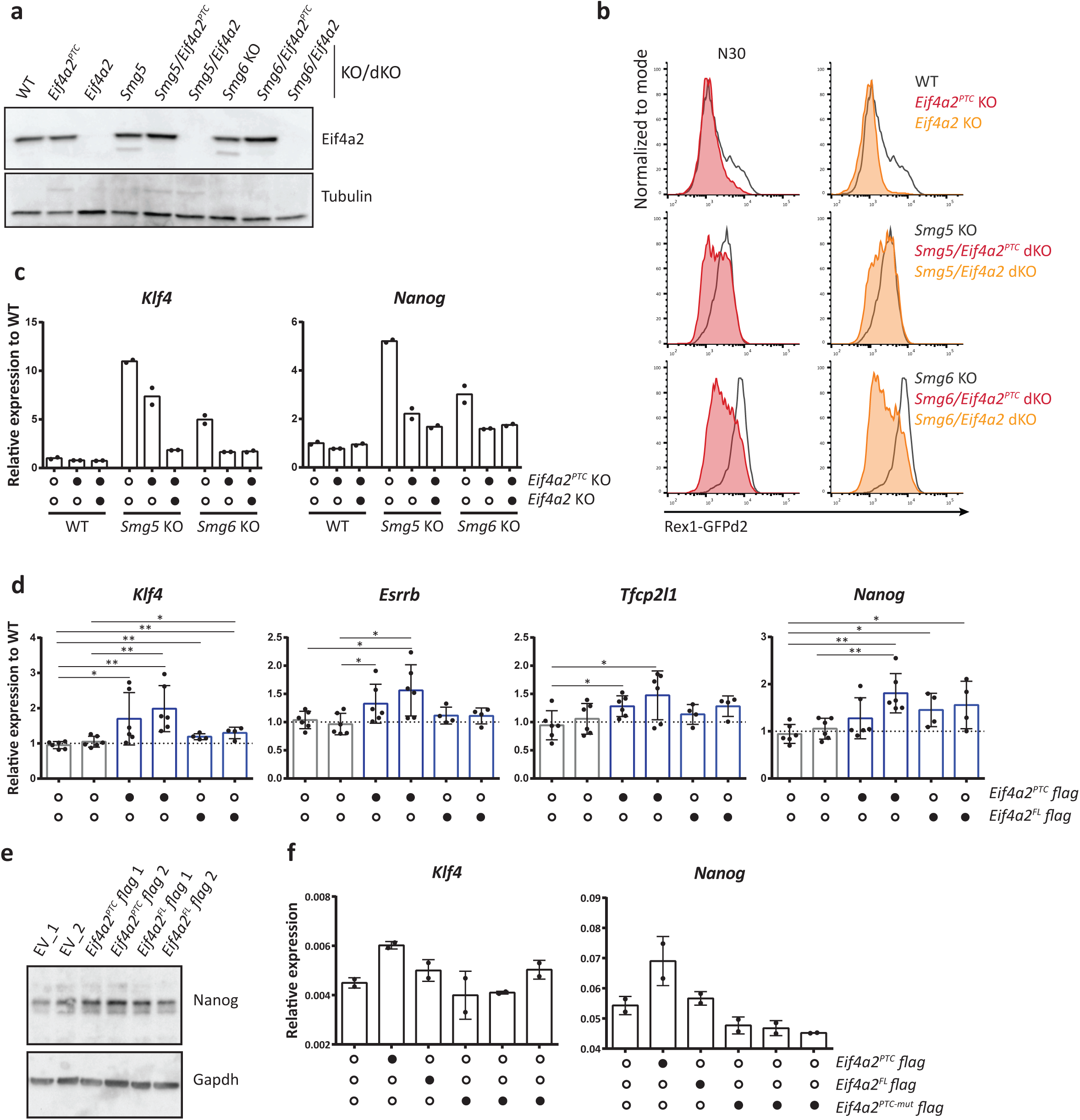
*Eif4a2* is causative for defects in exit from naïve pluripotency in NMD-deficient ESCs. **a**, Western blot analysis of Eif4a2 in the indicated cell lines. Tubulin was used as a loading control. **b**, Rex1-GFPd2 analysis at N30 in NMD KO and NMD/Eif4a2 dKO ESCs. Grey indicates FACS-profiles of WT cells, red indicates profiles of *Eif4a2^PTC^* deficient (*Eif4a2^PTC^* and NMD/Eif4a2^PTC^) cells and orange indicates profiles of *Eif4a2* deficient (*Eif4a2* KO and NMD/Eif4a2 dKO) cells. **c**, Representative experiment for the expression of indicated genes at N30 in NMD KO and NMD/Eif4a2 dKO cells. Mean of technical replicates plotted for each cell line. Expression was normalized first to *L32* and then to WT cells. **d**, Expression of the indicated genes at N30 in *Eif4a2* overexpressing and WT cells. Mean and SD are plotted for each cell line, n=3 biological replicates. Expression was normalized first to housekeeping genes (*actin* or *L32*). The expression level in WT cells was set to 1. Unpaired t-test was used to calculate p-values (* p < 0.05, ** p < 0.005). **e**, Western blot analysis for Nanog in *Eif4a2* overexpressing and WT cells at N24. Gapdh was used as loading control. **f**, Expression of the indicated genes at N16 in *Eif4a2* overexpressing and WT cells. Mean and SD of technical replicates are plotted for each cell line.

To assess the specificity of *Eif4a2* depletion in restoring differentiation potential in an NMD-deficient ESC background, we compared differentiation potential between *Tcf7l1* KO and *Eif4a2/Tcf7l1 dKO* ESCs. *Tcf7l1* encodes for a HMG box transcription factor and is one of the strongest drivers of the exit from naïve pluripotency*. Tcf7l1* KO ESCs showed defects in exit from naïve pluripotency on par with NMD KOs, but exhibited an NMD-independent, Gsk3 inhibition-like phenotype (Extended Data Fig. 5c) ^4^. *Tcf7l1* KO cells were insensitive to co-depletion of *Eif4a2* (Extended Data Fig. 5d), suggesting that Eif4a2 is a specific genetic interactor of NMD in regulating differentiation.

To test whether *Eif4a2* upregulation is sufficient to cause a differentiation delay, we overexpressed flag-tagged *Eif4a2^FL^* or *Eif4a2^PTC^* in WT ESCs (Extended Data Fig. 5e). At N24, *Eif4a2^PTC^* overexpressing cells showed increased transcript levels for *Klf4, Esrrb, Tfcp2l1* and *Nanog* and increased protein levels for Nanog at N24 (Fig. 5d,e). Cells overexpressing *Eif4a2^FL^* exhibited a weaker effect and showed mild upregulation only of *Nanog*. This accords with increase of Eif4a2^PTC^ rather than *Eif4a2^FL^* having a major impact on the exit from naïve pluripotency. Notably, the differentiation delays elicited by increasing expression levels of *Eif4a2* isoforms did not reach the intensity observed in NMD KO ESCs, suggesting that factors in addition to *Eif4a2* contribute to the failure to properly shut down naïve pluripotency in NMD KO ESCs. However, double depletion experiments show that deregulation of Eif4a2 is the major cause for exit from naïve pluripotency defects.

To test whether upregulation of *Eif4a2^PTC^* mRNA or Eif4a2^PTC^ protein was responsible for the delay in differentiation of NMD KO ESCs, we introduced a frameshift mutation in the first exon of *Eif4a2^PTC^* (Eif4a2^PTC-mut^) (Extended Data Fig. 5f). This mutation does not allow the generation of a functional Eif4a2 protein, abrogating in-frame coding potential. We overexpressed either *Eif4a2^PTC^* or *Eif4a2^PTC-mut^* in WT ESCs (Extended Data Fig. 5g). At N16, cells overexpressing *Eif4a2^PTC^* showed upregulation of *Nanog* and *Klf4* mRNA levels, while cells overexpressing *Eif4a2^PTC-mut^* did not (Fig. 5f). This suggests that upregulation of the truncated protein encoded by *Eif4a2^PTC^* is responsible for the differentiation delay observed in NMD KO cells. In summary, these data show that Eif4a2 is the NMD target causative for the observed differentiation delays in Smg-factor deficient ES cells.

### Eif4a2-mediated differentiation delay is caused by PTC isoform-dependent regulation of translation

Since Eif4a2 functions as a translation initiation factor, we tested whether its upregulation in NMD KO cells resulted in increased translation rates. To this end we performed radioactive S^35^ and O-propargyl-puromycin (OPP) based translation assays and detected an increase in translation rates in NMD-deficient cells in 2i and at N12 (Fig. 6a-c and Extended Data Fig. 6a). The increase in translation rates observed in NMD KO ESCs was similar in magnitude to that observed for *Tsc2* KOs, in which deregulated mTORC1 activity leads to an increase in translation ^32^. Full Eif4a2 KOs, but also specific depletion of the PTC isoform resulted in mild reductions of translation rates. Eif4a2^FL^ and Eif4a2^PTC^ depletion in NMD KO cells reduced translation rates to WT levels, showing that increased translation rates in NMD KO cells are dependent on the PTC isoform of Eif4a2 (Fig. 6b,c).

**Fig. 6.**
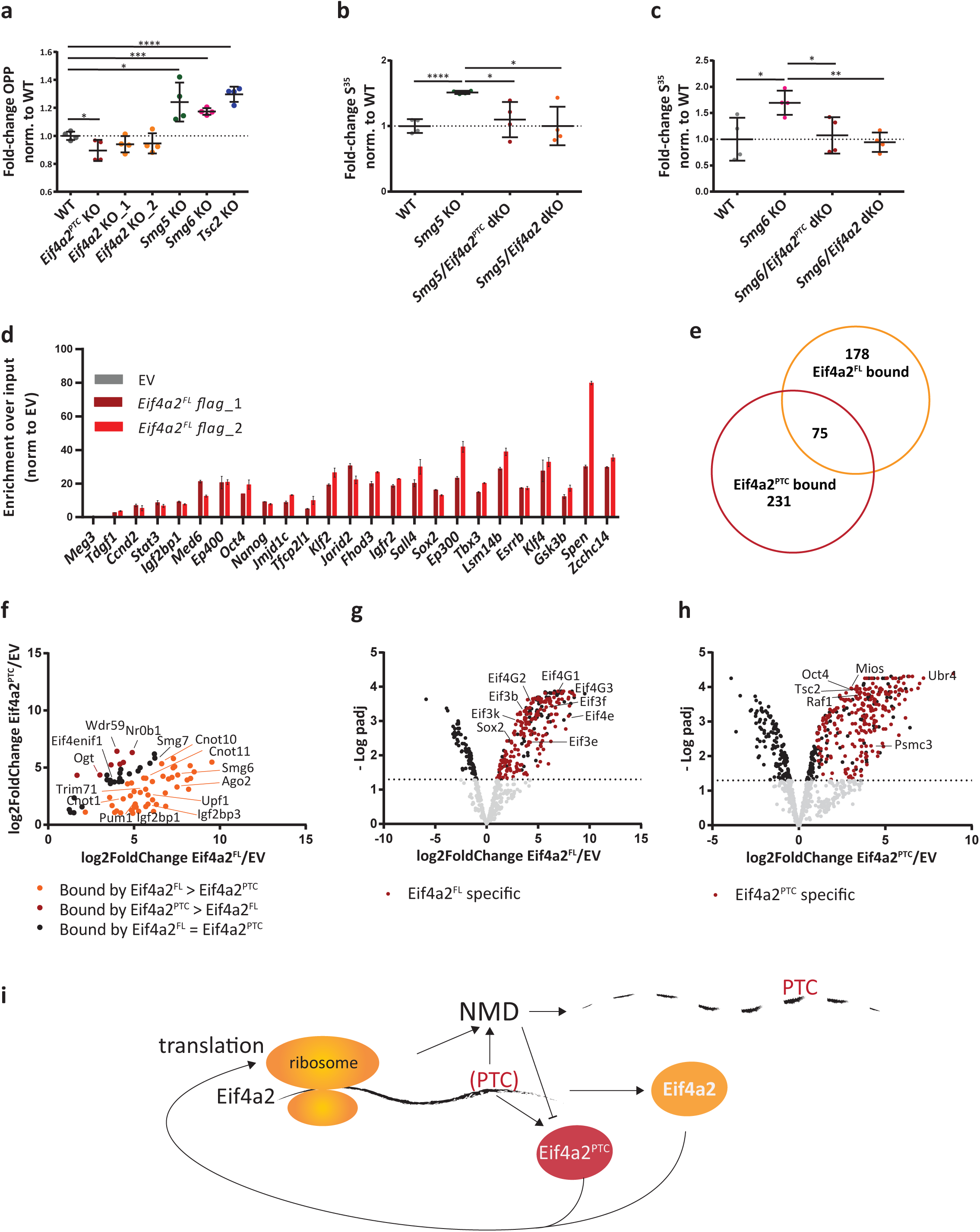
Eif4a2-mediated differentiation delay is caused by PTC isoform-dependent regulation of translation. **a**, OPP incorporation in NMD KO, *Eif4a2* KO, *Eif4a2^PTC^* KO, *Tsc2* KO and WT cells. Cells after 12 h (N12) of differentiation were pulsed for 20 s with OPP, followed by a 10 min chase. Mean and SD are plotted, n=4 biological replicates. Unpaired t-test was used to calculate p-values (**** p < 0.0001, *** p = 0.0001, * p < 0.05) **b**, S^35^ incorporation in *Smg5* KO, *Smg5/Eif4a2^PTC^* dKO, *Smg5/Eif4a2* dKO relative to WT. Cells in ES-DMEM-2i were pulsed with 40 µCi of S^35^ for 30 min before harvesting. Mean and SD are plotted; n=4 biological replicates. Unpaired t-test was used to calculate p-values (**** p < 0.0001, * p ≤ 0.05). **c**, S^35^ incorporation in *Smg6* KO, *Smg6/Eif4a2^PTC^* dKO, *Smg6/Eif4a2* dKO relative to WT. Experiment was performed as in 6b. Mean and SD are plotted, n=4 biological replicates. Unpaired t-test was used to calculate p-values (** p = 0.002, * p ≤ 0.05). **d**, Transcripts enriched by Eif4a2^FL^ RIP, detected by RT-qPCR. The non-protein-coding gene *Meg3* was used as negative control. Values were normalized to empty vector (EV)-RIP and to input. Mean and SD are plotted. Biological duplicates of *Eif4a2^FL^* RIP-samples are shown separately in shades of red. **e**, Significantly bound proteins (log2fold enrichment over EV ≥ 1; p-adj. ≤ 0.05) to Eif4a2^FL^ and Eif4a2^PTC^ identified by co-IP followed by mass-spec analysis. **f**, Comparison of enrichmentlevels of proteins significantly bound by both Eif4a2^FL^ and Eif4a2^PTC^ (75 proteins, see Figure 6e). **g**, Volcano plot showing proteins detected in mass-spectrometry after Eif4a2^FL^ immunoprecipitation. Grey dots represent detected proteins, black dots represent significantly enriched or depleted proteins and red dots represent proteins specifically enriched only in Eif4a2^FL^ co-IP (178 proteins, see Figure 6e). **h**, Volcano plot showing proteins detected in mass-spectrometry after Eif4a2^FL^ immunoprecipitation. Grey dots represent detected proteins, black dots represent significantly enriched or depleted proteins and red dots represent proteins specifically enriched only in Eif4a2^PTC^ co-IP (231 proteins, see Figure 6e). **i**, Summary model of NMD regulation indicating a proposed feedback circuit in which translation triggers NMD and NMD in turn controls transcript levels of a key translation initiation factor.

To identify transcripts whose translation was potentially most affected by Eif4a2, we performed RIP-Seq ^33^, using *Eif4a2* KO ESCs engineered to only express either flag-tagged Eif4a2^PTC^ or Eif4a2^FL^ (Extended Data Fig. 6b). As expected for a translation factor, noncoding transcripts like *Meg3* were depleted in flag-Eif4a2^FL^ RIP samples, indicating the specificity of the assay. Out of 9,812 expressed transcripts (CPM > 1 in all input samples), we detected 362 transcripts significantly bound by Eif4a2 (Extended Data Table 4). Several naïve pluripotency (*Esrrb, Tbx3, and Klf4*) and exit-factor (*Jarid2*, *Alg13* and *Fbxw7)* ^6^ encoding transcripts were enriched for Eif4a2 binding, as confirmed by RIP-qPCR (Fig. 6d and Extended Data Fig 6c). RIP-seq of flag-Eif4a2^PTC^ resulted in only weak enrichments, possibly owing to the unstable nature of the Eif4a2^PTC^ protein. RIP-RT-qPCR could not detect binding of Eif4a2^PTC^ to Eif4a2^FL^ targets identified above (Extended Data Fig. 6d).

To understand the interaction landscape of both *Eif4a2* isoforms, we used KO cells expressing either flag-tagged Eif4a2^PTC^ or Eif4a2^FL^ (Extended Data Fig. 6b). We also over-expressed Eif4a2^PTC^ in WT ESCs to identify potential Eif4a2^FL^-dependent interactions of Eif4a2^PTC^. We then performed co-IP followed by mass-spectrometry to identify interaction partners of the two different isoforms. To compensate for the highly unstable nature of Eif4a2^PTC^, we performed sample preparation after short-time treatment with the irreversible proteasome inhibitor epoxomicin ^34^. Thereby, we identified 254 and 306 proteins specifically bound by Eif4a2^FL^ or Eif4a2^PTC^ over flag-only control IPs, respectively (Extended Data Table 5 and Fig. 6e).

Among these, 75 proteins precipitated with both isoforms. Twenty-one of these showed near equal interactions between the Eif4a2 isoforms, including the NMD factor Smg7 and the translation initiation factor transporter Eif4enif1 (Extended Data Table 5), suggesting some preservation of normal interaction profiles of the PTC-specific isoform. Of the remainder, 47 showed stronger binding by Eif4a2^FL^. Among these were several negative regulators of mRNA stability and regulation (e.g., Ago2, Igfbp1 and Igfbp3, Pum1, Trim71, Cnot). We further detected physical association of Eif4a2^FL^ with several NMD components, such as Upf1 and Smg6, consistent with previous reports ^35^. Together, this indicates an intricate link between Eif4a2 and the mRNA destabilization machinery. We detected seven factors that showed stronger co-precipitation with the PTC-isoform. These include the pluripotency TF Nr0b1 (Dax1) ^8^, the pluripotency regulator Ogt ^36^ and the GATOR complex member and mTOR regulator Wdr59 (Fig. 6f).

Among the 178 proteins specifically bound to Eif4a2^FL^ and not to the PTC isoform were various translation initiation factors including all Eif4G isoforms and several components of the Eif3 complex (Fig. 6g and Extended Data Table 5). This highlights the integration of Eif4a^FL^ (but not Eif4a2^PTC^) into a functional translation initiation complex. Eif4a1 was not detected, indicating a mutually exclusive presence in the complex. Notably, Sox2 interacted with Eif4a2^FL^ (Fig. 6g), indicating a potentially functional interaction of core pluripotency regulators and the translation initiation machinery.

Among the 231 proteins specifically associated with the PTC isoform and not the full-length protein, we detected the Fgf/ERK pathway component Raf1, the mTORC1 regulator Tsc2 and the GATOR complex member Mios (Fig. 6h and Extended Data Table 5). Increased mTORC1 activity in NMD is evident by increased p-p70-S6K levels in NMD KO cells (Extended Data Fig. 6e). We further detected an interaction of Eif4a2^PTC^ with the key pluripotency factor Oct4 (*Pou5f1*).

Taken together, these results indicate that Eif4a2^FL^ is mainly involved in translation initiation, but shows significant interactions with a set of mRNA destabilizing proteins, including NMD-factors, CCR4-CNOT components and the RISC member Ago2. The unstable Eif4a2^PTC^ protein showed little association with translation initiation complex members, but still binds mRNA binding proteins, including the NMD factors Smg6 and Smg7.

In summary, we show that NMD controls transcript levels of hundreds of genes in ESCs and during early differentiation. This results in delayed exit from naïve pluripotency and a failure to shut down cell fate programmes in later cell fate decisions. Deregulation of a feedback circuit between NMD and translation initiation in NMD mutant cells, encoded in a PTC-containing isoform of Eif4a2, results in elevated translation rates, which are the major cause of NMD-associated differentiation defects in ESCs.

## DISCUSSION

Here we report a function of NMD in the dynamic regulation of a mammalian cell state transition, the exit from naïve pluripotency. Our results highlight a role for NMD far beyond purging PTC containing transcripts. We here show a major role in modulating gene expression profiles and also translation, suggesting NMD as important component of the cellular machinery maintaining cell identity and proper differentiation trajectories.

Without NMD function, increased levels of Eif4a2 and Eif4a2^PTC^ disrupt normal differentiation kinetics and lead to an increase in translation levels, including those of naïve pluripotency TFs, which impedes developmental transition. Basal leaky expression of differentiation-associated transcripts was reduced already in 2i. Consistently, NMD depletion leads to increased similarities between ESCs and the naïve *in vivo* epiblast ^6^. Therefore, NMD acts already in self-renewal conditions to prune the differentiation programme. Despite exhibiting delayed downregulation of the pluripotency TF-network, NMD KO EBs eventually initiate formative and primed expression programmes. However, subsequent downregulation of key markers of these states, normally observed around d4 to d6 of EB differentiation, does not occur. This suggests a general role for NMD in dynamically shaping and fine-tuning transcript abundance to facilitate rapid gene regulatory network remodeling during cell fate decisions.

Differentiation defects across *Smg5*, *Smg6* and *Smg7* KOs scale with deregulation of known NMD target genes and levels of phospho-Upf1, suggesting that NMD defects directly translate to differentiation delays. The striking difference in NMD target regulation and extent of differentiation delays between *Smg5* and *Smg7* KOs (Smg5 >> Smg7) together with the strong synergistic effects after co-depletion are difficult to reconcile with the proposed obligatory heterodimer dependence of these two factors ^24, 26^. We show that a division of labor between Smg5 and Smg7 is causative for the different effects of depletion in the respective KOs. Smg factors have a dual role in triggering RNA degradation and in mediating Upf1 dephosphorylation; pUpf1 increase in all three Smg KOs accords with such a role of Smg factors in PP2A recruitment ^25, 26^. Our data suggest that Smg7 acts as the main adapter or recruiter for pUpf1. However, by itself it is unable to efficiently dephosphorylate Upf1, as evidenced by high pUpf1 levels in *Smg5* KO ESCs, where Smg7 binding to pUpf1 is unaffected. We propose that Smg7 binding to Upf1 without subsequent Upf1 dephosphorylation results in jamming of the dephosphorylation cycle, and ensuing stalling of the mRNA degradation circuit. Aberrant recruitment of Smg6 to mRNA targets already bound by Smg7 and pUpf1 cannot properly restore NMD function in Smg5 KOs. Smg5 alone is unable to bind pUpf1 in our assays. This suggests that the main role of Smg5 lies in its dephosphorylation activity, but recruitment to its substrate is Smg7-dependent. This is consistent with the strongest increase of pUpf1 levels and consequently the strongest differentiation delays in *Smg5* KO ESCs.

Absence of *Smg6* causes a strong differentiation delay. This suggests that the exonucleolytic pathway cannot completely compensate for loss of the endonucleolytic activity in *Smg6* KOs. *Smg7* KO leads to very weak phenotypes and low levels of NMD deregulation. Our data support a model in which, in the absence of Smg7, Smg5 cannot be recruited to pUpf1-bound mRNAs, enabling Smg6 to recognize these potential targets and initiate their degradation. Thereby Smg6 can largely compensate for loss of Smg7, resulting in only mild NMD defects in Smg7 KO cells. However, Smg6 is unable to compensate for the complete absence of the exonucleolytic branch of NMD, suggested by the apparent loss of viability in a *Smg5/7* dKO situation. This indicates a function of Smg5 and/or Smg7 in the endonucleolytic decay pathway. Our data are consistent with a function of Smg5 or Smg7 in Upf1 dephosphorylation, which is a key step also in the endonucleolytic Smg6-mediated mRNA decay axis. Despite the weak defects in single *Smg7* KO cells, facultative essentiality for *Smg7* is revealed in the absence of either *Smg5* or *Smg6*. Accordingly, only *Smg5/Smg6* dKO ESCs could be established, showing that Smg7 is sufficient to sustain minimal NMD activity required for survival. This suggests that Smg7 can either interact with potentially novel interactors in the absence of its normal interaction partners, or that Smg7 acts as a jack of all trades, fulfilling minimal roles in target recognition, recruitment of the degradation machinery and Upf1 dephosphorylation. Taken together, the mode of action described above explains both the graded pUpf1 levels as well as graded impact on NMD of Smg5, Smg6 and Smg7 KOs.

A combination of RNA-seq and SLAM-seq based half-life analyses allowed us to compile a list of high-confidence NMD targets during the exit from pluripotency. Our data show partial redundancy and highly overlapping target gene sets between NMD downstream effectors in ESCs, consistent with previous results showing shared NMD targets between Smg6 and Smg7 ^19^. However, while target mRNA identity appears identical, the amplitude of mRNA deregulation scales with both the phenotype and the increase in pUpf1 levels.

NMD targets showing a graded upregulation pattern (Smg5 > Smg6 > Smg7) include the translation initiation factor *Eif4a2*. In contrast to previous reports, *c-myc* was not identified as an NMD target and showed no relevance for the differentiation defect in our experiments ^13^. We believe that our use of state-of-the-art ESC culture conditions facilitated the identification of differentiation-relevant NMD targets, that are obscured in heterogeneous FCS/LIF culture conditions.

In mammals, two homologs of the Eif4a helicase, Eif4a1 and Eif4a2, can be integrated into the eIF4F complex ^30, 31^. In our datasets only Eif4a2, and not Eif4a1, was found to be regulated by NMD. In addition to encoding for a full-length protein, the *Eif4a2* locus also produces a distinct PTC-containing isoform. In the truncated Eif4a2^PTC^ isoform 45 aa of the C-terminal helicase domain are missing, without affecting the ATP binding domain. Transcripts of both isoforms showed increased half-lives in NMD KO cells and reacted strongly to translation inhibition, a hallmark of bona fide NMD targets.

Control of protein synthesis rates is fundamental for maintenance of self-renewal and differentiation ^37, 38^ and for maintaining an ESC-specific chromatin state ^39, 40^. ESCs require a downregulation of translation rates and ribosome biogenesis to successfully differentiate ^40, 41^. We detected a marked increase in translation rates in NMD KO ESCs, which is not solely an effect of increased mRNA abundance due to increased mRNA stability, but directly dependent on increased Eif4a2 levels. This is evident from double-depletion experiments, in which *Eif4a2* depletion in NMD KOs reduces translation and partly rescues differentiation kinetics without restoring NMD function.

Increased protein levels of the Eif4a2^PTC^ isoform and not the full-length protein disrupt the normal translation programme and the exit from naïve pluripotency. This became evident in NMD KO cells in which the PTC exon was excised. In these cells, differentiation kinetics were restored despite increased expression of Eif4a2^FL^. Furthermore, only Eif4a2^PTC^ overexpression elicits clear differentiation delays. The function of Eif4a2^PTC^ appears dependent on the presence of Eif4a2^FL^ because overexpression of Eif4a2^PTC^ alone in Eif4a2 KO ESCs has no detectable impact on differentiation. This constitutes evidence that the translation initiation potential of Eif4a2 can be increased by its PTC isoform, which in turn requires the full-length protein to exert its function.

Eif4a2^PTC^ protein is detectably expressed in ESCs, but its physiological role remains to be determined. The observed interaction between Eif4a2^FL^ and Eif4a2^PTC^ with key pluripotency TFs is of special note. This interaction with the pluripotency circuitry remains to be explored, but could provide a direct link between translation initiation and cell fate control. Such a function accords with recent reports of Sox2 binding to the translation initiation factor transporter Eif4enif1, and Sox2 binding to RNA in mouse ESCs ^44, 45^.

NMD is triggered during translation. NMD-mediated regulation of Eif4a2 provides a link back from NMD to translation initiation, thereby establishing a feedback circuit (Fig. 6i). We propose that such a mechanism can be used to balance NMD activation with translational activity. We propose Eif4a2^PTC^ to act as a rheostat: Its upregulation upon NMD dysfunction increases translation initiation activity, which in turn directly increases the chances of triggering NMD in successive rounds of Eif4a2-initiated translation. In line with this hypothesis, our data confirm a reported physical interaction between Eif4a2 and Smg factors ^35^. Underscoring the importance of the Eif4a2^PTC^ mediated feedback loop to translation, the PTC-containing exon shows an unexpectedly high level of evolutionary conservation in various mammalian species, including human.

In conclusion, we show that NMD is required for proper restructuring of GRNs during differentiation by targeting multiple transcripts, most prominently the translation initiation factor Eif4a2. Our results place NMD as a central player in shaping transcriptomes to maintain proper cellular identity. We provide evidence for an intricate feedback circuit between NMD and translation. NMD susceptibility is hardwired in an evolutionarily conserved fashion into the *Eif4a2* gene structure and sequence. This direct link between NMD and translation initiation will need to be considered when studying NMD associated phenotypes. Our results are also relevant for improving therapeutic interventions and provide a rational for choosing the appropriate Smg-factor in pharmacological approaches to inhibit NMD activity, depending on the desired strength of NMD inactivation ^15^.

## AUTHOR CONTRIBUTIONS

EG performed experiments and analyzed data. RS, MR, FTT, LH and MG performed bioinformatic analysis, supervised by AB, AvH and CB. MH, JR, SS and LS performed experiments. CW and CS analyzed teratoma samples. AC, KFL and AP supported translation measurements. VH and SLA performed half-life measurements. EG, AS and ML designed experiments. ML supervised the study and wrote the manuscript together with EG with feedback from all authors.

## ACKNOWLEDGEMENTS

We thank Thomas Sauer and Johanna Stranner at the Max Perutz Labs FACS facility for expert support, Markus Hartl and the Massspec facility for mass-spec analysis and Irmgard Fischer for help with microscopy. NGS was performed at the VBCF. This work was supported by the Austrian Science Fund (FWF; P31334). Martin Leeb is a WWTF VRG group leader (VRG14-006) and was supported by an FWF Schroedinger return fellowship.

---

**Extended Data Fig. 1.**
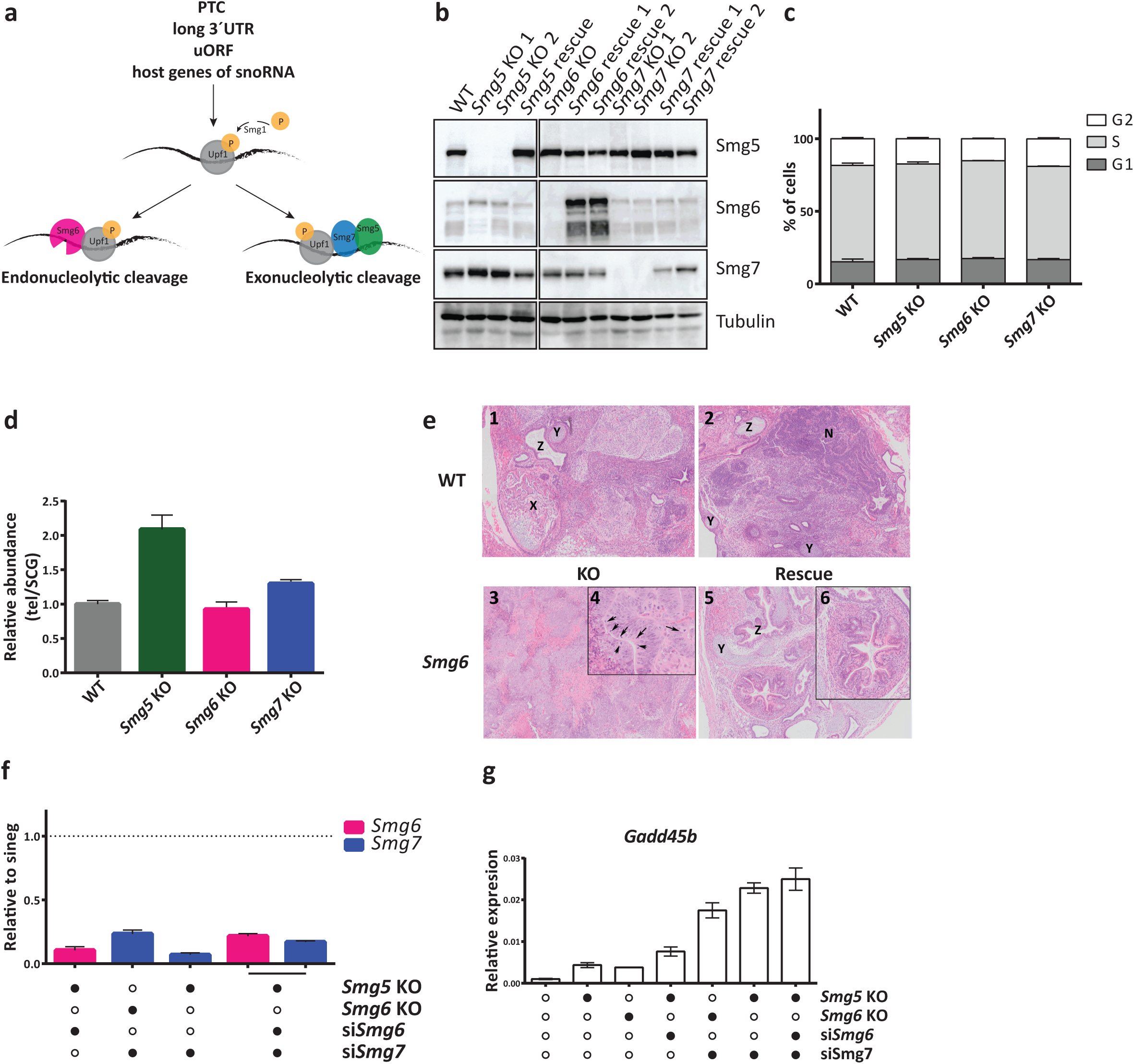
Hierarchy of defects in exit from naïve pluripotency in NMD-deficient ESCs. **a**, Illustration of the endo- and exonucleolytic branches of NMD. **b**, Western blot analysis in WT, NMD KO and NMD rescue ESCs using the indicated antibodies. Tubulin was used as loading control. **c**, Cell cycle analysis of NMD KO and WT ESCs. Propidium iodide was used to stain the cells and the Watson Pragmatic model was used to calculate the percentage of cells in each cell cycle phase. Mean and SEM are plotted for each cell cycle phase, n=3 biological replicates. **d**, Telomere length calculation in NMD KO and WT ESCs. Mean and SD of technical replicates are plotted for each cell line. Results were normalized to the 36B4 single copy locus. **e**, Teratomas derived from WT, *Smg6* KO and *Smg6* rescue ESCs. **1-2**: areas of WT-derived teratoma showing well differentiated tissues of mesodermal (e.g., X enchondral ossification; Y cartilage), endodermal (e.g., Z), and ectodermal (neuronal rosettes; N) origin. **3**: Area of *Smg6* KO-derived teratomas showing abundant poorly differentiated neuronal tissue; poorly differentiated endodermal and mesodermal tissues are present at lower abundancy (not shown). Abundant mitotic figures present in neuronal tissue (arrows) demonstrate high proliferative capacity (**4**). **5**: Area of *Smg6* rescue-derived teratomas showing examples of well differentiated endodermal (Z) and mesodermal (Y: cartilage) tissues. Higher power magnification shows an endodermal duct with different, well developed epithelial cell types (**6**). **f**, qPCR analysis confirming the siRNA-mediated knockdown of NMD components. Mean and SD of technical replicates are plotted for each cell line. Expression was normalized first to *L32*. Expression levels after sineg transfection were set to 1. **g**, Expression levels of *Gadd45b* in the indicated cells lines. Expression was normalized to *Actin*. Mean and SD of technical replicates are plotted for each cell line.

**Extended Data Fig. 2.**
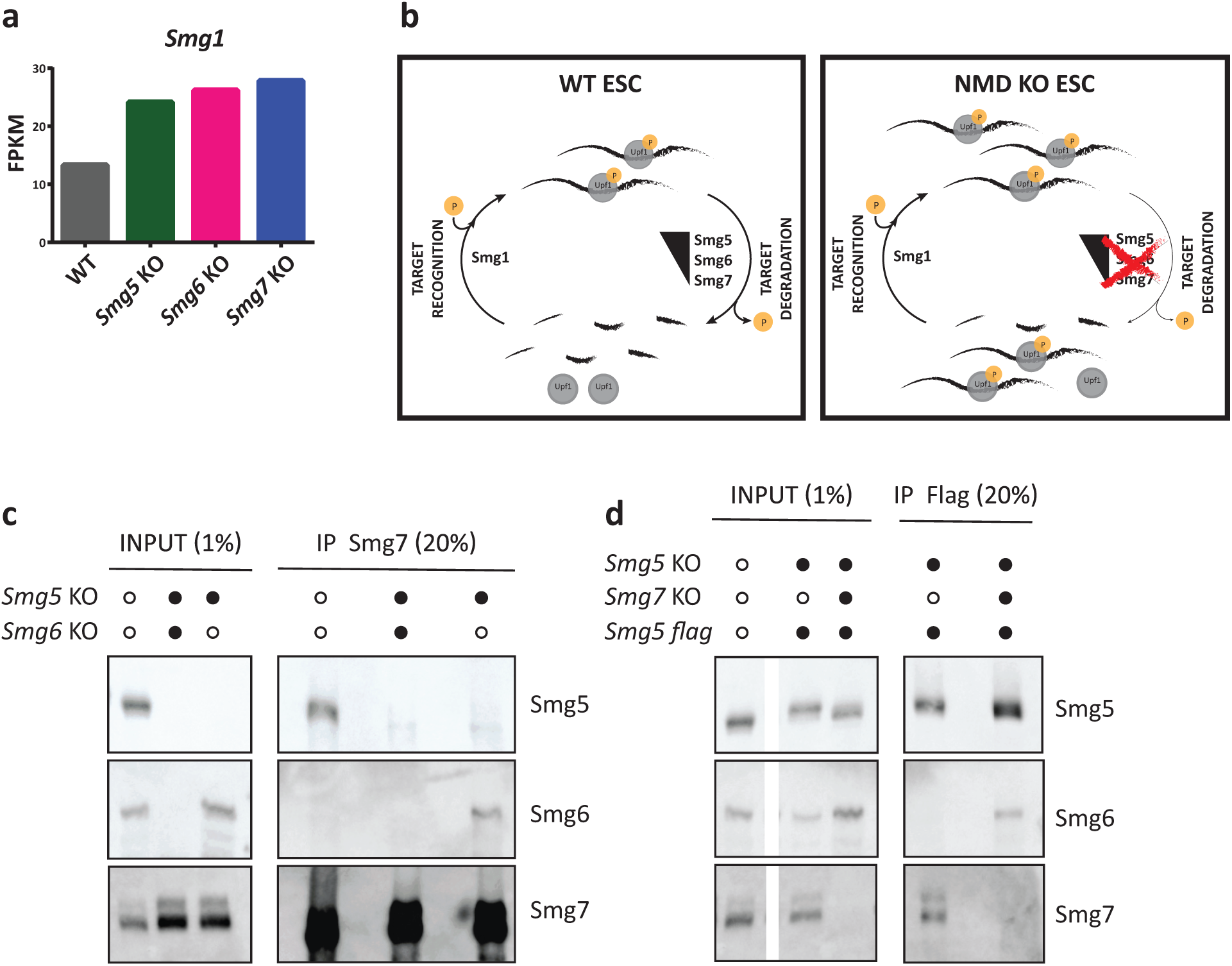
Smg7 is necessary and sufficient for pUpf1 binding, independently of Smg5. **a**, Expression levels (FPKM) of *Smg1* derived from RNA-seq in WT and NMD KO ESCs in 2i. Mean values between biological duplicates are plotted. **b**, Schematic representation of phosphorylation and dephosphorylation cycle of Upf1 in WT and NMD KO ESCs. **c**, Western blot of Smg7 co-IP in the indicated cell lines without cyanase treatment. Antibodies used are indicated in the figure. **d**, Western blot analysis of Smg5 co-IP (flag antibody) in the indicated cell lines without cyanase treatment. Antibodies used are indicated in the figure. **c** and **d** are parallel experiments to those shown in Figure 2b and c.

**Extended Data Fig. 3.**
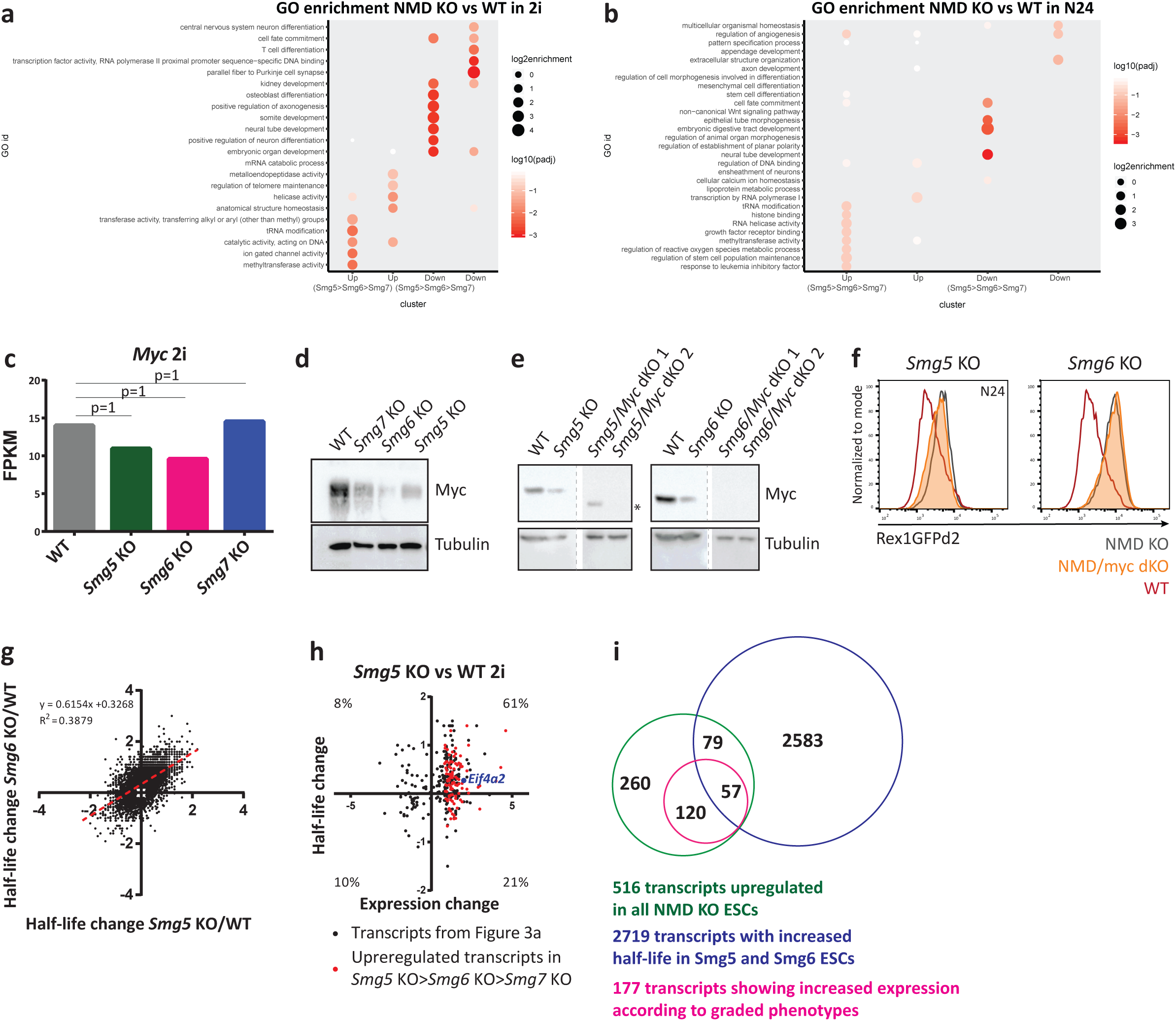
Integrating transcriptome-wide gene expression with mRNA half-life analyses identifies relevant NMD targets during the exit from naïve pluripotency. **a**, Significantly enriched GO terms of differentially expressed transcripts in NMD KOs in 2i. **b**, Similar to **(a)** for differentially expressed transcripts at N24. **c**, Expression levels of *c-myc* in NMD KO and WT ESCs in RNA-seq. Mean values between biological duplicates are plotted. p-adj. values are indicated above each bar. **d**, Western blot analysis in NMD KO and WT ESCs using the antibodies indicated. Tubulin was used as loading control. **e**, Western blot analysis confirming absence of myc protein in NMD/myc dKO ESCs. Antibodies are indicated in the figure. Tubulin was used as loading control. * band corresponds to truncated protein. **f**, Rex1-GFPd2 FACS profile at N24 in NMD KO and NMD/myc dKO cells. FACS profile of NMD KO cells is grey, orange profile shows NMD/myc dKO cells and red profile shows WT. **g**, Global analysis of NMD-induced half-life change in *Smg5* KO and *Smg6* KO ESCs (all transcripts with a half-life change ≥ |0.2| in Smg5 or Smg6 KO ESCs are plotted. Half-life change was calculated for 7,451 transcripts. Slope and R2 values are shown in the figure. **h**, Transcriptome-wide analysis of changes in mRNA expression and half-lives in *Smg5* KO. Half-life changes could be calculated for 321 out of the 516 transcripts upregulated in NMD KO ESCs (Figure 3a). 182 of the 516 transcripts showed a concomitant increase in half-life (half-life change ≥ 0.2). Transcripts behaving according to the graded phenotype are depicted in red. **i**, Venn diagram showing 516 transcripts upregulated in all NMD KO ESCs intersected with 177 transcripts showing graded increase in expression in NMD factor KOs (Smg5 > Smg6 > Smg7) and 2,719 transcripts with increased half-lives in both *Smg5* and *Smg6* KO ESCs.

**Extended Data Fig. 4.**
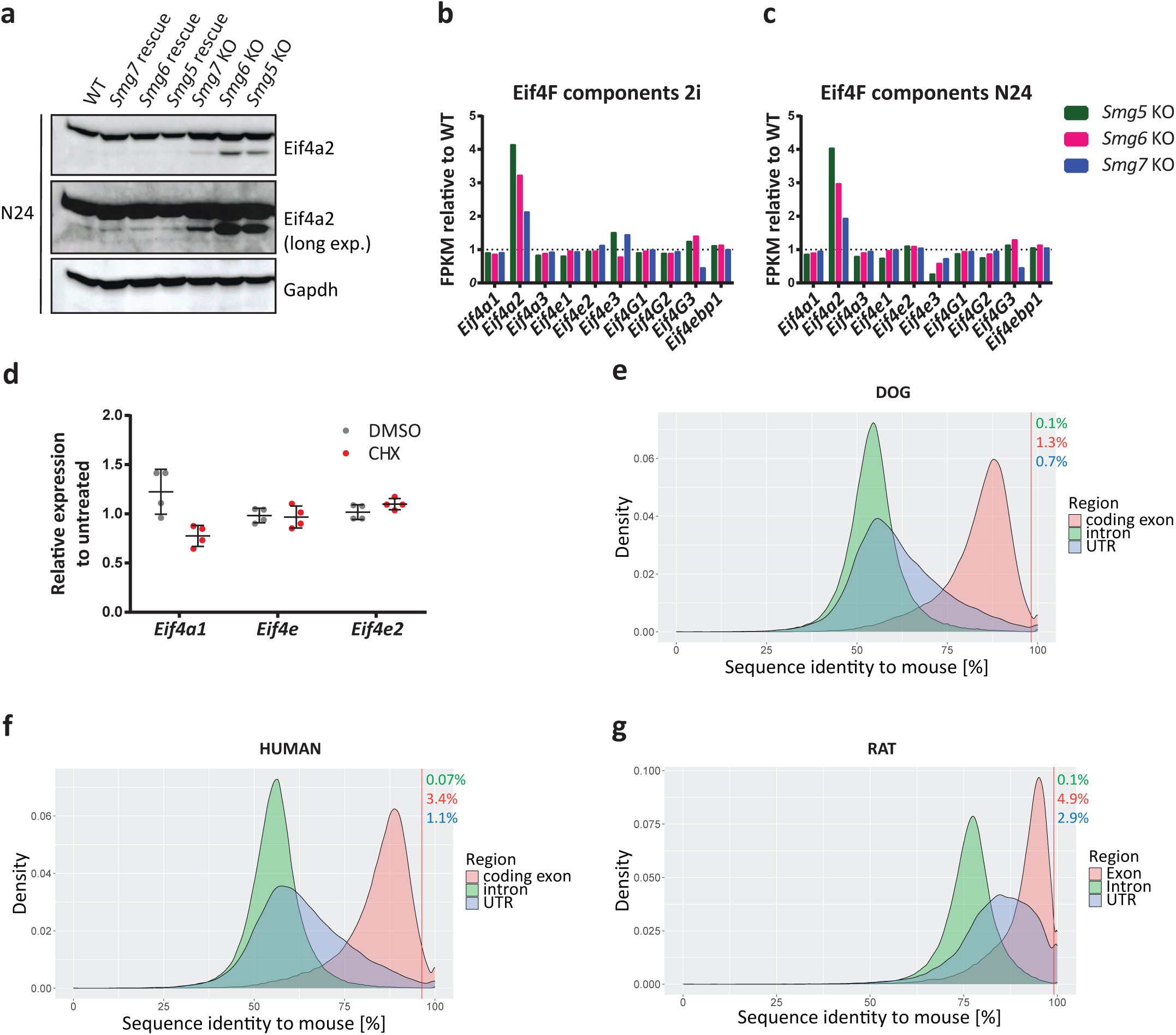
Eif4a2 is a bona-fide NMD target in ESCs. **a**, Western blot analysis for Eif4a2 protein levels in NMD KO, NMD rescue and WT cells at N24. Gapdh was used as loading control. **b**, Expression of Eif4F complex components and regulators in NMD KO ESCs relative to WT ESCs in self-renewal (2i). FPKM values from RNA-seq analysis are shown, see Figure 3a. **c**, Expression of Eif4F complex components and regulators in NMD KO cells relative to WT cells at N24. Values from RNA-seq analysis, see Figure 3c. **d**, Expression levels of the indicated genes after treatment with CHX or DMSO in WT ESCs, assayed by RT-qPCR. Mean and SD are plotted, n=2 biological replicates. **e-g**, Densities of the distributions of the sequence identities of indicated regions of the mouse transcriptome compared to dog (**e**), human (**f**) and rat (**g**). Red line indicates identity of PTC-containing *Eif4a2* exon. Percentage of genomic sequences with higher conservation than PTC-containing exon of *Eif4a2* are indicated in the graphs.

**Extended Data Fig. 5.**
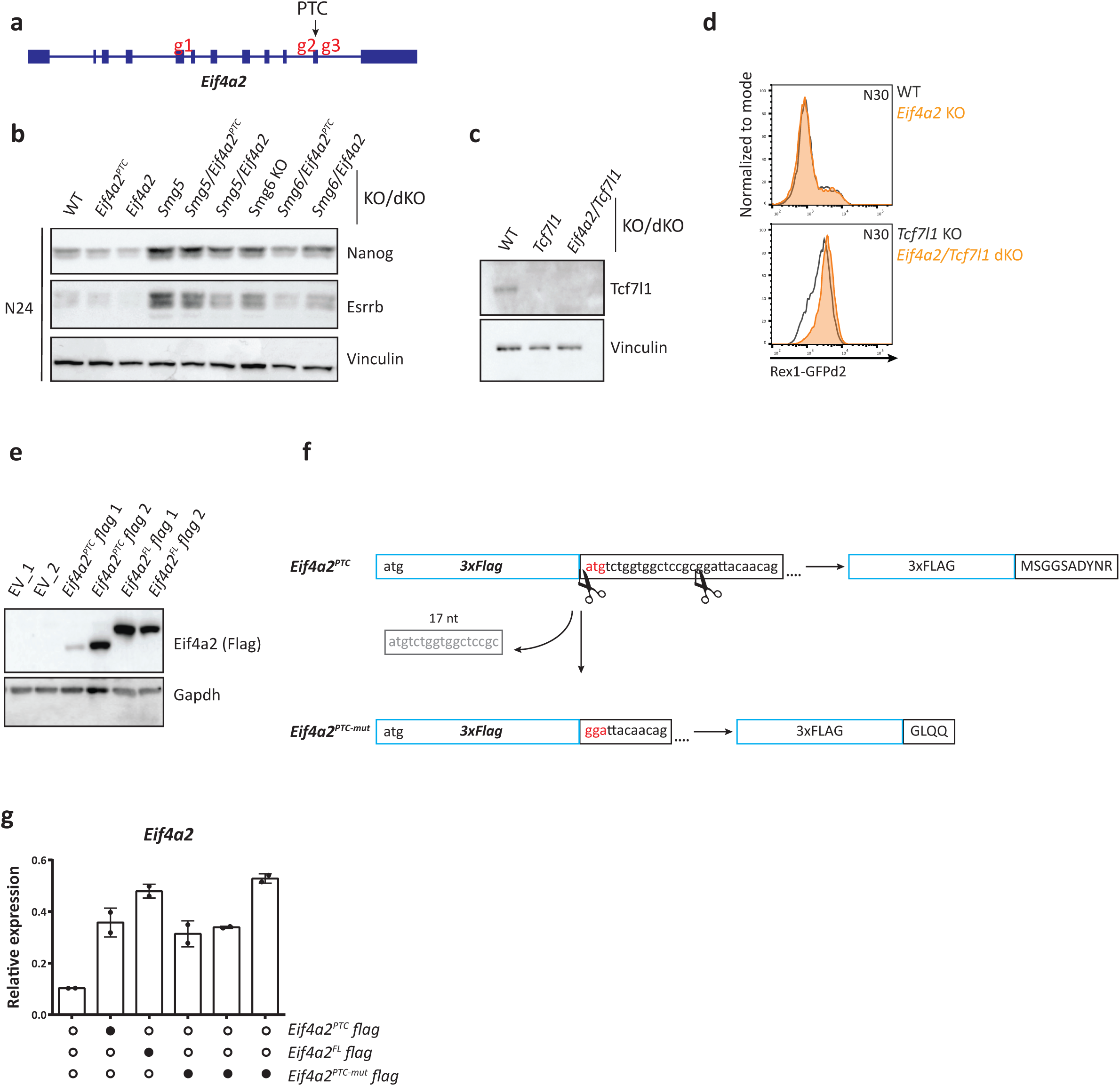
*Eif4a2* is causative for defects in exit from naïve pluripotency in NMD-deficient ESCs. **a**, Schematic view of the Eif*4a2* gene. gRNAs (g1 to g3) used to generate *Eif4a2* and *Eif4a2^PTC^* depletions are indicated. g1 and g3 were used together to delete all *Eif4a2* isoforms. g2 and g3 were used together to delete the *Eif4a2^PTC^* isoform. **b**, Western blot analysis at N24 in the annotated cell lines for pluripotency markers Esrrb and Nanog. Vinculin was used as loading control. **c**, Western blot analysis for Tcf7l1 expression in the indicated cell lines. Vinculin was used as loading control. **d**, Rex1-GFPd2 analysis at N30 in WT (grey), *Eif4a2* KO (orange), *Tcf7l1* KO (grey) and *Eif4a2/Tcf7l1* dKO (orange) cells. **e**, Western blot analysis of Eif4a2 (flag antibody) in *Eif4a2* overexpressing Eif4a2 KO cells. EV transfection shows no signal. Gapdh was used as loading control. **f**, Schematic illustration of the mutation introduced into the *Eif4a2^PTC^* isoform. **g**, Expression of *Eif4a2* at N16 in *Eif4a2* overexpressing pools and WT cells. Mean and SD of technical replicates are plotted for each cell line.

**Extended Data Fig. 6.**
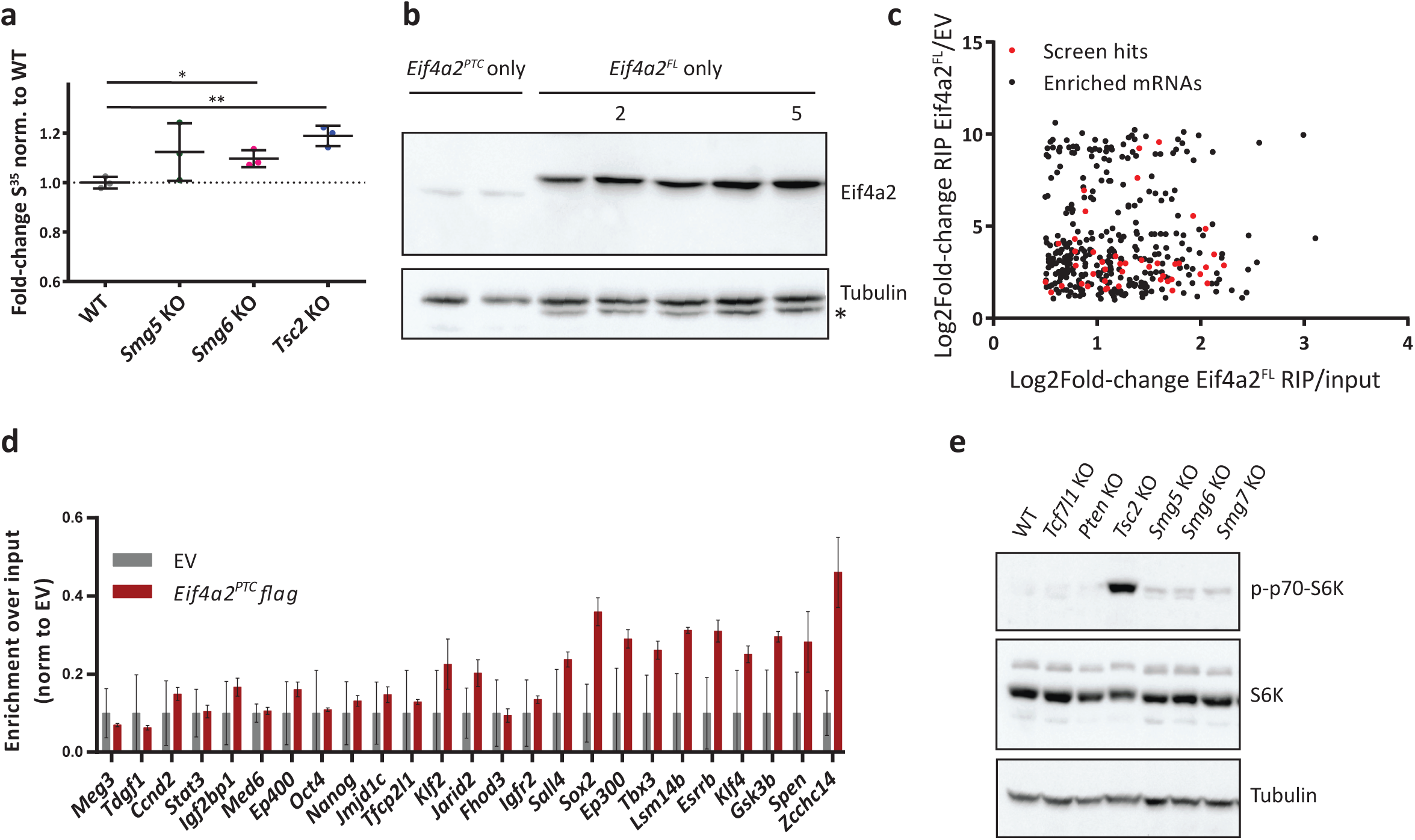
Eif4a2-mediated differentiation delay is caused by PTC isoform-dependent regulation of translation. **a**, S^35^ incorporation levels in NMD KO, *Tsc2* KO and WT ESCs. Experiment was performed independent from, but as described for Figure 6b. Mean and SD are plotted, n=2 biological replicates. Unpaired t-test was used to calculate p-values (* p-value = 0.02, ** p-value = 0.002). **b**, Western blot analysis for Eif4a2 in *Eif4a2* KO cells in which Eif4a2^PTC^ or Eif4a2^FL^ were re-expressed from transposon vectors. Tubulin was used as a loading control. For Eif4a2^FL^, only clones 2 and 5 were used for subsequent analysis. * corresponds to Eif4a2 band due to insufficient stripping. **c**, Significantly enriched transcripts in Eif4a2^FL^ RIP-Seq (log2fold change Eif4a2^FL^ RIP/input ≥ 0.5, p ≤ 0.05 and log2fold change Eif4a2^FL^ RIP/EV RIP ≥ 1, p ≤ 0.05). Transcripts of genes with shown function as drivers of the exit from naïve pluripotency ^6^ are depicted in red. **d**, Transcripts levels of the indicated genes after Eif4a2^PTC^ RIP-qPCR. The non-protein coding gene *Meg3* was used as negative control. Values were first normalized to EV RIP and then to input. Mean and SD are plotted, n=2 biological replicates. **e**, Western blot analysis in the indicated cells line in 2i for mTOR targets. Antibodies used are indicated in the figure. Tubulin was used as loading control.

## Material and Methods

### Cloning

gRNAs were designed using http://crisprscan.org. Annealed oligos (Extended Data Table 5) were cloned, using a BsaI site, in a gRNA expressing vector (Addgene #41824) ^46 6^. For generating rescue cell lines, the coding sequence of the gene of interest was amplified by PCR and cloned into a pCAG-3xFLAG-empty-pgk-hph vector ^5^ using a BamHI site. The sequence of Eif4a2^FL^ was *in vitro* synthetized (IDT) and then cloned into the pCAG-3xFLAG-empty-pgk-hph vector. For the mutagenesis of Eif4a2^PTC^ the pCAG-3xFLAG-Eif4a2^PTC^-pgk-hph vector was cut using a BamHI and a SacII site. This generated an Eif4a2^PTC^ fragment that was missing the first 17 nucleotides, thus generating an out of frame truncated protein product. The mutated insert was blunt ended and re-cloned into the pCAG-3xFLAG-empty-pgk-hph vector cut with SmaI. The correct insertion was verified for all the constructs by restriction digest and Sanger sequencing with the 3xFlag_seq primer (Extended Data Table 6).

### Teratoma assay

Paraffin-embedded teratoma tissue blocks were cut on a rotary microtome RM2255 (Leica). The 3 µm sections were then stained for hematoxylin and eosin in the automated slide stainer Gemini AS (Histocom) and mounted in Eukitt. Slides were scanned on a VS120 (Olympus) slide scanner.

### Cell culture

Mouse embryonic stem cells (ESCs) containing a Rex1-GFPd2-IRES-BSD Cas9 (Rex1-GFPd2) ^4, 23^ were used as parental cell line for all the knockout cells generated in this study.

ESCs were cultured on gelatin (Sigma-Aldrich, G1890) coated plates in DMEM high-glucose (Sigma-Aldrich, D5671) supplemented with 15% FBS (Gibco, 10270-106), 2 mM L-Glutamine (Sigma-Aldrich, G7513), 0.1 mM NEAA (Sigma-Aldrich, M7145), 1 mM Sodium Pyruvate (Sigma-Aldrich, S8636), 10 µg/ml penicillin-streptomycin (Sigma-Aldrich, P4333), 55 µM β-mercaptoethanol (Fisher-Scientific, 21985-023), 2i (1.5 μM CHIR99021 and 0.5 μM PD0325901) and 10 ng/ml LIF (batch tested, in-house) (ES DMEM-2i medium).

### Monolayer differentiation

For differentiation, ESCs were plated on gelatin-coated plates at a density of 1 × 10^4^ cells/cm^2^ in basal medium (N2B27), which is composed by 1:1 ratio of DMEM/F12 (Gibco, 21331020) and Neurobasal medium (Gibco, 21103049) supplemented with 0.5x N2 (homemade), 1x B-27 Serum-Free Supplement (Gibco, 17504-044), 2 mM L-Glutamine (Sigma-Aldrich, G7513), 0.1 mM NEAA (Sigma-Aldrich, M7145), 10 µg/ml penicillin-streptomycin (Sigma-Aldrich, P4333), 55 µM β-mercaptoethanol (Fisher-Scientific, 21985-023) and 2i (3 μM CHIR99021 and 1 μM PD0325901) (N2B27-2i). The following day 2i were withdrawn and cells were differentiated for the indicated time in N2B27.

### Commitment assay

For commitment assay, ESCs were plated on gelatin-coated plates in N2B27-2i medium at a density of 2 × 10^3^ cells/cm^2^. The following day 2i were withdrawn. After 48 h of differentiation medium was changed and selection for Rex1-GFPd2 expressing cells was carried out by adding 5 ng/ml blasticidin ((BSD) Gibco, R210-01) to the N2B27 medium. The following day medium was changed to N2B27-2i + BSD medium, and medium was refreshed every two days. Alkaline-phosphatase staining was performed after four days in N2B27-2i + BSD according to the manufacturer’s protocol (Sigma-Aldrich, 86R). In brief, cells were fixed using the Citrate-Acetone-Formaldehyde solution. Cells were then stained for 30 min at 4°C using a Sodium Nitrite-FRV-Alkaline-Naphthol AS-BI Alkaline solution. Plates were then imaged using an Olympus Cell-Sense microscope (OLYMPUS).

### EB differentiation assay

For embryoid body differentiation, 1 × 10^5^ cells were used and plated in ES DMEM without 2i and LIF as hanging drops. After two days cells were collected and plated in a 10 cm petri dish, and medium was changed every two days. Cells were harvested every two days and RNA was extracted using the ExtractME kit (LabConsulting, EM09).

### Generation of knockout cell lines

2 × 10^5^ cells were transfected in a gelatin-coated 6-well plate in ES DMEM-2i using Lipofectamine 2000 (Fisher Scientific, 11668-027). Two pairs of gRNA were used for each gene (1 µg each) together with 0.5 µg of pCAG-dsRed (see full list of gRNAs in Extended Table 6). After 48 h to enrich for transfected cells, dsRED/GFP double positive cells were sorted using a BD FACS Aria III. Sorted cells were plated at clonal density in ES DMEM-2i. One week after sorting 48 colonies were picked (96 in case of the Smg5/Smg6 dKO cells). Colonies were then trypsinized and half of the cell suspension was plated in a 96-well plate for expansion. The remaining cells were lysed and DNA was extracted for PCR genotyping ^6^.

### PCR genotyping

For DNA lysis half of a picked colony was pelleted in a PCR plate at 500g. After two PBS washes, cell pellets were boiled in water at 95°C for 5 min. 3 µg/ml of proteinase K was added after cooling down the plate and incubated at 65°C for 1 h. Reaction was inactivated for 10 min at 95°C. To genotype the KO cells we used a triple-primer strategy. One reverse primer specific for the deletion and one specific for the WT allele were used in combination with a common forward primer (Extended Data Table 6). PCR results were verified by Sanger sequencing and KO was confirmed by western-blot analysis. For genotyping PCR JumpStart RedTaq PCR master mix (Sigma-Aldrich) was used following the manufacturer’s protocol for cycling with an annealing temperature of 55°C and 35 cycles.

### RNAi assay

1.5 × 10^4^ cells/cm^2^ were transfected in a gelatin-coated 12-well plate in N2B27-2i using DharmaFECT 1 (Fisher Scientific, T-2001). For RNAi assays esiRNA for Eif4a2 (Sigma-Aldrich) and FlexiTube siRNAs for Smg6 and Smg7 (Qiagen) were used. For siRNAs 20 ng siRNAs/4 × 10^4^ cells were used, whereas for esiRNAs 200 ng/6 × 10^4^ cells were used. The following day, after two PBS washes, medium was changed to N2B27. Cells were harvested at the indicated time for either flow cytometry to determine Rex1-GFPd2 activity (see flow cytometry) or for RNA extraction (see RNA analysis). Primers used for qPCR are annotated in Extended Data Table 6.

### Cell cycle analysis

ESCs were plated in gelatin-coated plates in ES DMEM-2i at a density of 1 × 10^4^ cells/cm^2^. After two days cells were harvested and fixed overnight in 70% ethanol. Cells were then incubated with 100 μg/ml RNase A solution (Qiagen, 19101). 50 mg/liter Propidium Iodide solution (Sigma-Aldrich, P4170) was then added to the cells. Cell cycle profiles were recorded on the LSRFortessa flow cytometer (BD bioscience).

### Flow Cytometry

Cells were harvested using 0.25% trypsin/EDTA and trypsin was neutralized using ES DMEM. Rex1-GFPd2 was measured with LSRFortessa flow cytometer (BD bioscience). High-throughput-measurements were acquired using a 96-well plate HTS unit on the LSRFortessa flow cytometer. Data analysis was performed using flowJo software (BD bioscience).

### NMD inhibition assay

ESCs were plated in gelatin-coated plates in ES DMEM-2i at a density of 2 × 10^4^ cells/cm^2^. The following day cells were treated for 8 h with either ES DMEM-2i + DMSO or ES DMEM-2i + CHX (100 ug/ml (Sigma-Aldrich, 01810)). RNA was then extracted using ExtractME kit (LabConsulting, EM09) according to the manufacturer’s protocol.

### Telomere quantification

For telomere quantification genomic DNA was extracted from cells using the Puregene core Kit A (Qiagen). DNA was quantified using PicoGreen Assay for dsDNA (Fisher Scientific, P11496) on a NanoDrop 3300 Fluorospectrometer (Fisher Scientific). PCR reactions were performed on a CFX384 Touch Real-Time PCR Detection System (BioRad) using telomere specific primers and primers for the single-copy gene 36B4 (Extended Data Table 6). For each primer pair, a standard curve was created with known amounts of DNA to determine primer efficiency. The telomere signal was normalized with the single-copy gene.

### RNA analysis

RNA was extracted using the ExtractMe kit (LabConsulting, EM09) according to the manufactureŕs protocol. cDNA was retrotranscribed from 0.3 µg to 1 µg using the SensiFAST cDNA Synthesis Kit (Bioline, BIO-65054). Real-time PCR was performed on the CFX384 Touch Real-time PCR Detection System (Bio-Rad) using the Sensifast SYBR No Rox-Kit (Bioline, BIO-98020). Expression levels were normalized to either L32 or Actin, as indicated in the figure legends (see full list of primers used for qPCR In Extended Data Table 6). Results are shown as mean and standard deviation.

### Protein analysis

Proteins were extracted using RIPA buffer (Sigma-Aldrich, 20-188) supplemented with Complete Mini EDTA-free Protease inhibitor cocktail (Roche, 04693159001) and PhosSTOP (Phosphatase Inhibitor Cocktail (Roche, 04906845001)). 5% milk was used for blocking. Primary antibodies were incubated overnight at 4°C. Washes were performed using PBS-T (Sigma-Aldrich, P4417). Secondary antibodies were incubated for 1 h at room-temperature. Primary antibodies were used at a dilution of 1:250 for anti-Smg5 (Abcam, ab33033, rabbit), 1:1000 for anti-Smg6 (Abcam, ab87539, rabbit), 1:2000 for anti Smg7 (NovusBio, NBP1-22967, rabbit), 1:1000 for anti-Phosho-Upf1 (Millipore, 07-1016, rabbit), 1:1000 for anti-Upf1 (D15G6) (Cell Signaling, 12040S, rabbit), 1:1000 for anti-flag M2 (Sigma-Aldrich, F1804, mouse), 1:1000 for anti-c-Myc (D84C12) (Cell Signaling, 5605T, rabbit), 1:1000 for anti-Tcf7l1 (Fisher Scientific, PA5-40327, rabbit), 1:1000 for anti-Eif4a2 (Abcam, ab31218, rabbit), 1:1000 for anti-Eif4a1 (Cell Signaling, 2490T, rabbit), 1:1000 for anti-Nanog (NovusBio, NB100-58842, rabbit), 1:1000 for anti-Esrrb (RND systems, PP-H6705-00, mouse), 1:1000 for anti-p-p70-S6K (Thr389) (108D2) (Cell Signaling, 9234T, rabbit), 1:1000 for anti-S6K (49D7) (Cell Signaling, 2708T, rabbit), 1:5000 for anti-Tubulin (Sigma-Aldrich, T8203, mouse), 1:5000 for anti-Gapdh (Sigma-Aldrich, G8795, mouse) and 1:5000 for anti-Vinculin (E1E9V) (Cell-Signaling, 13901T, rabbit). Secondary antibodies were used at a dilution of 1:10000 for anti-rabbit IgG (Amersham, NA934), 1:15000 for goat anti-mouse IgG (Santa Cruz, sc-2064). Chemiluminescence signal from antibody binding was detected using ECL Select detection kit (GE healthcare, GERPN2235) with a ChemiDoc system (BioRad).

### Immunoprecipitation

Cells were plated in ES DMEM-2i, after two days they were harvested in IP lysis buffer (10 mM Tris Base, 10 mM NaCl, 2 mM EDTA, 0,5% Triton X-100) supplemented with Complete Mini EDTA-free Protease inhibitor cocktail (Roche, 04693159001) and PhosSTOP (Phosphatase Inhibitor Cocktail (Roche, 04906845001)). 1 mg of lysate was used for the immunoprecipitation. Briefly, Dynabeads (Fisher Scientific, 10004D) were coated with 5 µg of anti-flag M2 antibody (Sigma-Aldrich, F1804, mouse) for 1 h. In case of RNA-free IP, lysates were treated with 50 U/ml Cyanase (Süd-Laborbedarf SLG, CY1000) and 0.5 ul/ml of 2 M MnSO4 and incubated on ice for 10 min. Lysates were then cleared out by centrifugation at 16000 *g* for 15 min at 4°C, 1% of each lysate was kept as input. NaCl was added to a final concentration of 150 mM. Dynabeads were washed three times with wash buffer (137 mM NaCl, 20 mM Tris Base, 0,5% (v/v) Tergitol-type NP-40) and one time with lysis buffer and then were incubated with the lysates for 3 h at 4°C. Dynabeads were washed three times with wash buffer supplemented with Complete Mini EDTA-free Protease inhibitor cocktail (Roche, 04693159001) and PhosSTOP (Phosphatase Inhibitor Cocktail (Roche, 04906845001)). Samples were eluted in 2X sample buffer.

For the immunoprecipitation coupled with mass-spec cells were plated in ES DMEM-2i, the following day they were treated with 1 µM epoxomicin (Gentaur, 607-A2606) for 3 h. Cells were harvested and lysed in IP lysis buffer. 1 mg of lysate was used for the immunoprecipitation. Immunoprecipitation protocol was followed (see above) with the addition of the cross-linking of the anti-flag M2 antibody to the Dynabeads using dimethyl pimelimidate (DMP) (Sigma-Aldrich, D-8388). Briefly, after antibody coupling, Dynabeads were washed three times with 200 mM Sodium Borate (pH=9) and then incubated for 30 min with DMP-Sodium Borate solution. They were washed with the following buffers: three times with 250 mM Tris (pH=8.0), two times with 100 mM glycine (pH=2), three times with TBS-T and one time with lysis buffer. Beads were then incubated with the lysates for 2 h at 4°C. Five washes were performed with wash buffer (137 mM NaCl, 20 mM Tris Base) and samples were submitted for mass-spec.

### Sample preparation for mass spectrometry analysis

Beads with cross-linked antibody were transferred to new tubes and resuspended in 30 µL of 2 M urea in 50 mM ammonium bicarbonate (ABC). Disulfide bonds were reduced with 10 mM dithiothreitol for 30 min at room temperature before adding 25 mM iodoacetamide and incubating for 15 min at room temperature in the dark. Remaining iodoacetamide was quenched by adding 5 mM DTT and the proteins were digested with 150 ng trypsin (Trypsin Gold, Promega) at room temperature for 90 min. The supernatant was transferred to a new tube, the beads were washed with another 30 µL of 2 M urea in 50 mM ABC and the wash combined with the supernatant. After diluting to 1 M urea with 50 mM ABC, additional 150 ng trypsin were added and incubated overnight at 37°C in the dark. The digest was stopped by addition of trifluoroacetic acid (TFA) to a final concentration of 0.5 %, and the peptides were desalted using C18 Stagetips ^47^. Peptides were separated on an Ultimate 3000 RSLC nano-flow chromatography system (Thermo-Fisher), using a pre-column for sample loading (Acclaim PepMap C18, 2 cm × 0.1 mm, 5 μm, Thermo-Fisher), and a C18 analytical column (Acclaim PepMap C18, 50 cm × 0.75 mm, 2 μm, Thermo-Fisher), applying a segmented linear gradient from 2% to 35% and finally 80% solvent B (80% acetonitrile, 0.1% formic acid; solvent A 0.1% formic acid) at a flow rate of 230 nL/min over 120 min. Eluting peptides were analyzed on a Q Exactive HF-X Orbitrap mass spectrometer (Thermo Fisher), which was coupled to the column with a customized nano-spray EASY-Spray ion-source (Thermo-Fisher) using coated emitter tips (New Objective).

### mRNA half-life measurement

ESCs were plated in N2B27-2i at a density of 20 × 10^4^ cells/cm^2^. The day after medium was changed to N2B27-2i + 100 µM 4SU (Carbosynth, NT06186) and they were cultured in this condition for 12 h. After a PBS wash, cells were incubated with N2B27-2i medium + 10 mM Uridine (Sigma-Aldrich, U6381) for 3 h. RNA extraction was carried out using Trizol (Fisher-Scientific, 10296-010) following the manufacturer’s instruction with the addition of 0.1 mM of DTT during the isopropanol precipitation. RNA was resuspended in 1 mM DTT. 5 µg of RNA were treated with 10 mM iodoacetamide (Sigma-Aldrich, I1149) followed by ethanol precipitation ^29^. 2 ng of RNA were used for library prep. Libraries were prepared using the QuantSeq 3′mRNA-seq Library Prep Kit for Illumina FWD (Lexogen, R3142) and were analyzed on a HiSeqV4 SR100.

### RNA-Immunoprecipitation

3 × 10^6^ cells were plated in ES-DMEM. After two days cells were harvested and lysed using the MAGNA-RIP kit (Millipore 17-700) according to the manufactureŕs protocol. Briefly, cells were lysed by a freeze and thaw cycle and lysates were stored at −80°C. Immunoprecipitations were performed at 4°C for 3 h using 5 µg of an anti-FLAG M2 antibody (Sigma-Aldrich, F1804, mouse). For RIP-Seq, libraries were prepared using the QuantSeq 3′mRNA-seq Library Prep Kit for Illumina FWD (Lexogen, R3142) and analyzed on a HiSeqV4 SR50. For RIP-qPCR, reverse transcription and qPCR were carried out as above. Relative binding to input and empty vector control was calculated. Error bars show the standard deviation between technical duplicates (Eif4a2^FL^ RIP) and between biological duplicates (Empty vector and Eif4a2^PTC^ RIP).

### Translation rates measurement with S^35^ incorporation

ESCs were cultured in ES DMEM and 33µCi S^35^ (Hartmann, IS-103) was added to the culture media for 30 min. Cells were lysed in RIPA buffer (Sigma-Aldrich, 20-188) supplemented with Complete Mini EDTA-free Protease inhibitor cocktail (Roche, 04693159001) as described above. Protein extracts were then spotted on a nitrocellulose membrane. Membranes were stained with Ponceau. The membranes were wrapped in saran wrap and exposed to a BAS Storage Phosphor Screen (GE Healthcare). After two days signal was acquired using a Typhoon scanner (GE Healthcare). Radioactive signal was quantified using Fiji. S^35^ signal was normalized to ponceau staining. Results are shown as mean and standard deviation.

### Translation rates measurement with OPP incorporation

ESCs were cultured in N2B27 2i medium, the following day 2i were withdrawn. After 12 h of differentiation the medium was changed to N2B27 supplemented with OPP (Fisher Scientific) for a 20 s pulse. Cells were then washed two times in N2B27 medium and then N2B27 medium was added for a 10 min chase. Cells were then harvested. Fixation, permeabilization and Click-IT reaction were performed using the Click-iT Plus OPP Alexa Fluor 647 Protein Synthesis Assay Kit (Fisher Scientific, C10458), according to the manufactureŕs protocol. Fluorescent signal was acquired with LSRFortessa flow cytometer (BD bioscience). Median OPP-647 signal was used for quantification. Results are shown as mean and standard deviation.

### Data analysis

#### RNA-seq differential analysis

RNA-seq samples and analysis used in this study were taken from Lackner and colleagues ^6^

#### GO enrichment analysis

GO analysis annotation were taken from the R package org.Mm.eg.db (Version 3.4.0). For the enrichment of GO Terms in the different gene lists we only considered terms with 5 to 500 genes assigned to them. Significance of the enrichment was determined using Fisher’s exact test with all expressed genes as background (Lackner et al., 2020). GO terms that did not differ in more than 5 genes were clustered and one representative term for each cluster was defined (using the R base function hclust on the L1-distance of the binary membership matrix). The representative term for each cluster was the term with the least total annotated genes. Multiple hypothesis testing was executed using the Benjamini-Hochberg method on all representative terms to calculate adjusted p-values.

#### mRNA half-lives calculation

QuantSeq data was analyzed using SLAM-DUNK (Version 0.2.4). For calculation of RNA half-lives, background (no 4SU treatment) was subtracted to T>C conversion rates of transcripts with CPM ≥2. Half-lives were calculated by single exponential fit to a decay model ^29^.

#### mRNA half-lives differential analysis

T>C conversion rates were used to model the half-life changes. Background (no 4SU treatment) was subtracted from T>C conversion rates and then T>C conversion rates of transcripts with CPM ≥2 were considered for the differential analysis. Differential analysis to quantify the change in T>C conversion rates was calculated with a beta regression model (betareg package, version 3.1-3) of the form: T>C conversion rates ∼ time * genotype. In brief, firstly the T>C conversion rates after 3 h of Uridine chase were normalized to the T>C conversion rates of the 4SU pulse. Secondly, the beta regression model quantified the difference between T>C conversion rates of WT and NMD KO cells after 3 h of Uridine chase. p-values were determined by a partial Wald test. To determine transcripts with a significant change in expression and half-life we considered the 516 upregulated transcripts from Figure 3a. We could calculate a half-life change for 250 out of 516 transcripts. To obtain direct targets, we postulated that upregulated transcripts should have a concomitant increase in half-life (half-life change ≥ 0.2). Finally, to select NMD targets relevant in ESCs out of these 136 transcripts we isolated the 57 that were upregulated in expression according to the graded phenotypes.

### Exon conservation

Pairwise whole genome alignments of mouse (GRCm38/mm10) against dog (canFam3), rat (RGSC6.0/rn6) and human (GRCh38/hg38) from UCSC (http://hgdownload.cse.ucsc.edu/downloads.html) were used for the analysis. With the Ensembl annotation of the mouse genome we extracted the respective regions of the genes (introns, exons and UTRs) and calculated their sequence identities compared to the other organism in the pairwise alignment as (number of matching nucleotide pairs in the alignment)*100/length(alignment). As gene region annotation can be ambiguous due to variations in transcript splicing, we used the following definitions. Everything that is annotated as coding sequence in any protein coding transcript we count as coding exon. UTR is everything that is annotated as UTR in any protein coding transcript except if it is also annotated as coding sequence in which case it is counted as coding exon. Regions not covered by the above definitions are counted as introns.

### RIP-Seq differential analysis

Quality control of the transcripts from input and RIP samples was performed using fastQC. Reads were trimmed using bbduk (Version 38.57). Reads were mapped to the mm10 mouse reference genome with STAR (Version 2.5.3a). Afterwards indexing was performed using samtool (Version 1.5) and reads in transcripts were counted with HTSeq-count (Version 0.11.2). Transcripts that had cpm more or equal to one in all the input samples were considered for the differential analysis. Differential analysis was performed using DESeq2 (Version 1.24.0). To identify significantly enriched transcripts in Eif4a2^FL^ RIP we considered only the 1,231 transcripts with cpm ≥30 in Eif4a2^FL^RIP and either log-fold-change Eif4a2^FL^ input / Eif4a2^FL^ RIP ≥ 1 and ≤-1 (p-value ≤0.05) or log-fold-change Eif4a2^FL^ RIP / EV RIP ≥ 1 and ≤-1 (p-value ≤0.05). Out of those we considered as enriched only the transcripts that had Eif4a2^FL^ input / Eif4a2^FL^ RIP ≥ 0.5 and log-fold-change Eif4a2^FL^ RIP / EV RIP ≥ 1 (362 transcripts).

### Mass spectrometry data acquisition and analysis

The mass spectrometer was operated in data-dependent acquisition mode (DDA), survey scans were obtained in a mass range of 375-1500 m/z with lock mass activated, at a resolution of 120k at 200 m/z and an AGC target value of 3E6. The 8 most intense ions were selected with an isolation width of 1.6 m/z, fragmented in the HCD cell at 28% collision energy and the spectra recorded for max. 250 ms at a target value of 1E5 and a resolution of 30k. Peptides with a charge of +1 or >+7 were excluded from fragmentation, the peptide match feature was set to preferred, the exclude isotope feature was enabled, and selected precursors were dynamically excluded from repeated sampling for 30 s.

Raw data were processed using the MaxQuant software package (version 1.6.0.16, ^48^) and the Uniprot mouse reference proteome (January 2019, www.uniprot.org), target sequences, as well as a database of most common contaminants. The search was performed with full trypsin specificity and a maximum of two missed cleavages at a protein and peptide spectrum match false discovery rate of 1%. Carbamidomethylation of cysteine residues were set as fixed, oxidation of methionine and N-terminal acetylation as variable modifications. For label-free quantification the “match between runs” feature and the LFQ function were activated - all other parameters were left at default.

MaxQuant output tables were further processed in R (R Core Team, 2018, https://www.R-project.org/). Reverse database identifications, contaminant proteins, protein groups identified only by a modified peptide, protein groups with less than three quantitative values in one experimental group, and protein groups with less than 2 razor peptides were removed for further analysis. Due to differences in overall contaminant levels between samples, LFQ values were re-normalized using the sample median of the “background” protein subset (as identified in controls). Missing values were replaced by randomly drawing data points from a normal distribution modeled on the whole dataset (data mean shifted by −1.8 standard deviations, width of distribution of 0.3 standard deviations). Differences between groups were statistically evaluated using the LIMMA package ^49^ at 5% FDR (Benjamini-Hochberg). The mass spectrometry proteomics data have been deposited to the ProteomeXchange Consortium via the PRIDE partner repository ^50^ under the following accession number PXD019588. NGS data have been deposited on GEO (https://www.ncbi.nlm.nih.gov/geo/query/acc.cgi?acc=GSE153457, reviewer token: uvglmauwnfufpkp)

